# The roles of aridification and sea level changes in the diversification and persistence of freshwater fish lineages

**DOI:** 10.1101/2020.01.27.922427

**Authors:** Sean J Buckley, Chris Brauer, Peter Unmack, Michael Hammer, Luciano B. Beheregaray

## Abstract

While the influence of Pleistocene climatic changes on divergence and speciation has been well-documented across the globe, complex spatial interactions between hydrology and eustatics over longer timeframes may also determine species evolutionary trajectories. Within the Australian continent, glacial cycles were not associated with changes in ice cover and instead largely resulted in fluctuations from moist to arid conditions across the landscape. Here, we investigate the role of hydrological and coastal topographic changes brought about by Plio-Pleistocene climatic changes on the biogeographic history of a small Australian freshwater fish, the southern pygmy perch *Nannoperca australis*. Using 7,958 ddRAD-seq (double digest restriction-site associated DNA) loci and 45,104 filtered SNPs, we combined phylogenetic, coalescent and species distribution analyses to investigate the relative roles of aridification, sea level and tectonics and their associated biogeographic changes across southeast Australia. Sea-level changes since the Pliocene and reduction or disappearance of large waterbodies throughout the Pleistocene were determining factors in strong divergence across the clade, including the initial formation and maintenance of a cryptic species, *N.* ‘flindersi’. Isolated climatic refugia and fragmentation due to lack of connected waterways maintained the identity and divergence of inter- and intraspecific lineages. Our historical findings suggest that predicted increases in aridification and sea level due to anthropogenic climate change might result in markedly different demographic impacts, both spatially and across different landscape types.

## INTRODUCTION

Dramatic changes in climate, hydrology and topography have long been recognised to have lasting impacts on the diversity, distribution and divergence of species and populations (Pelletier et al. 2015). Understanding the relationship between the historical environment and the genealogy of species remains critical for interpreting how contemporary climate change may impact on species currently and in the near future. Most notably, increasing aridification and rising sea-levels predicted by climate change projections call into question the adaptive capacity and resilience of organisms, especially those with poor dispersal potential and narrow ranges (Davis et al. 2013, Falkenmark 2013, Grummer et al. 2019). These effects are particularly exacerbated within regions of highly heterogeneous topography and climatic variation which can lead to diverse and multifaceted impacts on species (Guarnizo and Cannatella 2013, Graae et al. 2018). Applying broad-scale inferences about environmental changes to understand historical biogeography and biodiversity resilience in the future is further complicated by spatial variation in environmental factors that might impact on how within-species responses vary across their ranges (Razgour et al. 2019).

The relative role of Earth history events on the evolution and persistence of species is expected to vary across regions (Barton et al. 2013). Even a single major event may present multifaceted impacts on species evolution depending on how local or regional environments are shaped (e.g. sea level changes; Lambeck et al. 2012). For example, studies have highlighted the role of glacial refugia throughout Pleistocene glacial – interglacial cycles driving distributional shifts across the northern hemisphere, particularly within the Americas and Europe (Carnaval et al. 2009, Duckett et al. 2013, Pelletier et al. 2015). Although glacial ice expansion/retraction is unlikely to have affected much of the southern hemisphere throughout these cycles *per se* (Duckett et al. 2013, Lamb et al. 2019), secondary impacts such as intensifying aridity, sea-level changes and temperature shifts during glacial maxima have impacted on the evolution, distribution and persistence of southern hemisphere biotas (Williams et al. 2018, Ansari et al. 2019). Such secondary impacts likely shape the environment of different regions based on their local relevance, with eustatic changes having a larger influence on coastal or marine ecosystems while aridification played a stronger role further inland (Beheregaray et al. 2002, Pinceel et al. 2013). Additionally, environmental changes associated with tectonic shifts or the formation and submergence of land-bridges are mostly locally relevant and vary across landscapes. Thus, understanding the relative role of different environmental changes between regions is important in more accurately predicting species’ responses.

Inferences of phylogeographic responses to past environmental change relies upon a combination of genetic, spatial and modelling approaches. However, determining the relative role and chronology of past climatic events is difficult when resolution is low due to few genetic markers or limited model capability (Carstens et al. 2012, Cutter 2013, Nakhleh 2013). To this end, the development of next-generation sequencing allows for the collection of thousands of genetic markers which better capture the diverse array of demographic processes influenced by Earth history (Carstens et al. 2012, Edwards et al. 2016). In tandem, recent advancements in coalescent modelling, informed by more detailed information of geological and ecological history, have improved the ability to provide more nuanced inferences (Cutter 2013, Excoffier et al. 2013). This combination of greater data and more sophisticated modelling techniques provides the analytical framework to address questions about the spatial variance of species responses to climate change.

An ideal biogeographic setting to test hypotheses of spatial and temporal variation of Earth history on evolution is one including both inland and coastal regions. In this regard, the temperate southeast of Australia is well-suited given it has been influenced by aridification across the continent as well as by shifts in landmass attributed to eustatic changes (Faulks et al. 2010, McLaren and Wallace 2010, Chapple et al. 2011). This region is characterised by complex geography and geology, affected by a history of uplift, subsidence and volcanism (Unmack 2001). There is a strong gradient in temperature and precipitation across the region, with cooler and wetter climates towards the south. The region is subdivided by the Great Dividing Range, which runs parallel to the coastline from the top of Australia down to the southern coast. This range acts as a barrier that separates the inland Murray-Darling Basin (MDB) from coastal areas and is a key biogeographic feature of the region (Fig. 1b; Unmack 2001, Chapple et al. 2011).

**Figure 1:**
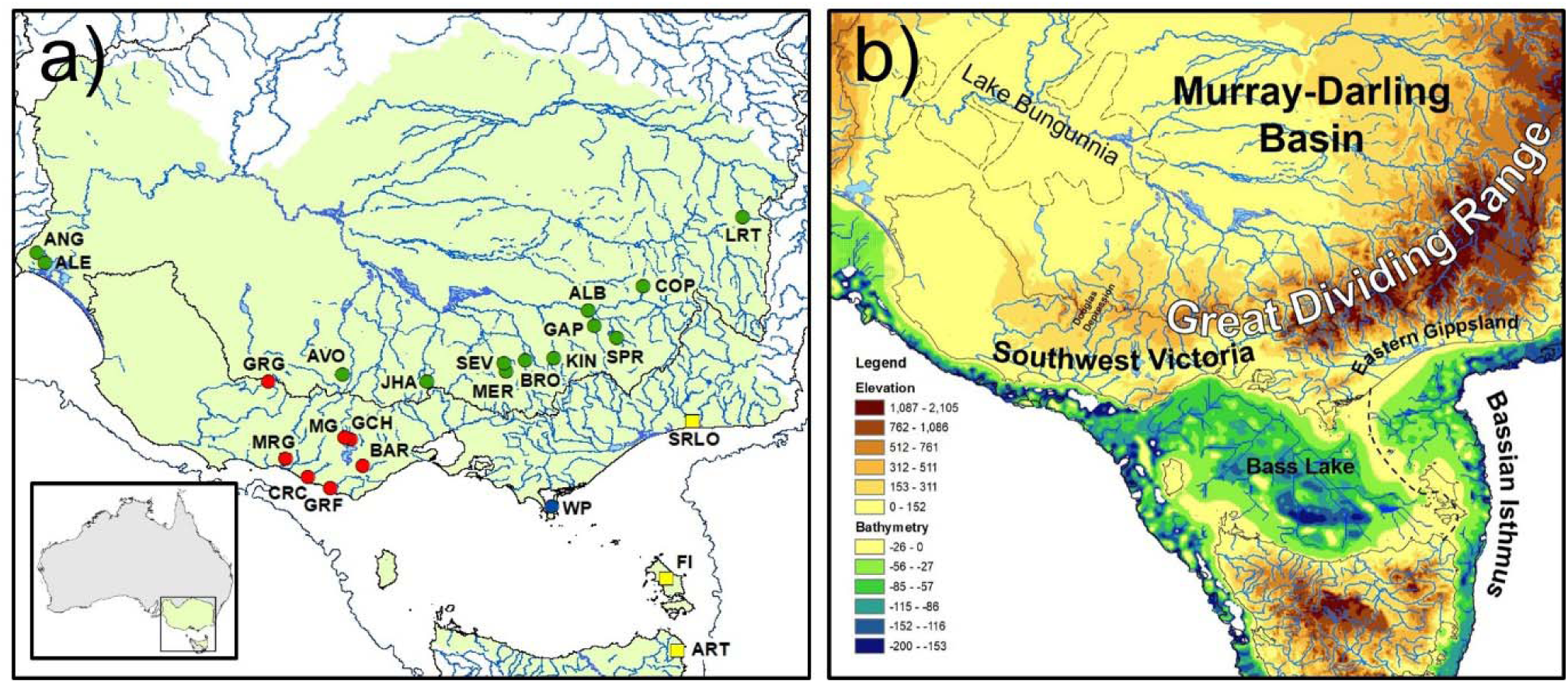
**a)** Distribution and sampling map for southern pygmy perch. Inset depicts extent of distribution within Australia. The shaded area denotes the putative distribution of the species, spanning multiple major basins (black lines). Colours denote major clades explored within coalescent models (refer to Results) whilst shapes denote ‘species’ (circles *= N. australis*; squares = *N.* ‘flindersi’). The extent of the continental shelf (−121m), which was exposed during glacial periods, is indicated in dark blue. **b)** Topographic map (including bathymetry) of southeast Australia, highlighting topographic heterogeneity and major biogeographic regions across the area. Solid black lines indicate major basin boundaries whilst the bold dashed line indicates the drainage divide across the Bassian Isthmus. The maximum extent of Lake Bungunnia (at 1.2 Ma) is also indicated with a narrow dashed line.

The MDB is one of the continent’s largest freshwater basins and a key water source for much of Australia. The MDB is hydroclimatically variable, with notable differences in hydrology and climate between headwaters and the terminal lowland lakes and wetlands near the Murray mouth (Pittock and Finlayson 2011). A major environmental change that reshaped the MDB over time was the formation and decline of the paleo megalake Bungunnia, which spanned 90,000 km^2^ across the lower section of the MDB at its largest size (McLaren et al. 2011, McLaren et al. 2012). Lake Bungunnia initially formed in the late Pliocene ∼3 Ma when tectonic uplift dammed the outlet of the ancestral Murray River. Its level fluctuated in accordance with glacial and interglacial climate cycles; interglacial periods caused the lake to reach overfilling where it resembled an open system of freshwater lakes (McLaren et al. 2012). Predictions of the volume of inflow required to maintain Lake Bungunnia at that size suggests 2 – 3 times higher rainfall than currently experienced was needed (Stephenson 1986, McLaren and Wallace 2010). Repeated overfilling of Lake Bungunnia during wetter climatic periods led to the eventual erosion of a new outlet to the west, resulting in the removal of the barrier and complete drainage of the inland lake system ∼700 Ka (McLaren and Wallace 2010). Lake Bungunnia has been suggested to be a relevant factor in the phylogeography of some terrestrial species (Cooper et al. 2000, Joseph et al. 2008, Kawakami et al. 2009, Ansari et al. 2019, Neal et al. 2019), and its fluctuation is likely an important biogeographic aspect of the region for local aquatic species (Waters et al. 2019).

These attributes contrast with the coastal habitats south of the MDB, where major changes in the environment are more associated with eustatic changes. One example is the formation of the Bass Strait which separated Tasmania from the mainland. Historically, Tasmania was connected to the mainland through a land bridge known as the Bassian Isthmus: central to this landmass was a large freshwater lake known as the Bass Lake (Blom and Alsop 1988, Porter-Smith et al. 2012). As sea levels rose during each interglacial of the Pleistocene, the Bassian Isthmus was inundated. As a result, the land bridge narrowed into an eastern corridor prior to its submergence, the remnants of which can be seen in the Flinders and Cape Barren Islands. The isolation of Tasmania and the formation of the Bassian Isthmus as the sole terrestrial connection are well documented drivers of biogeographic patterns for a variety of both terrestrial, marine and freshwater taxa (Schultz et al. 2008, Waters 2008). This combination of relevant Earth history factors across southeast Australia, and the potentially interactive nature of these events, provides an ideal scenario to investigate the relative role of different past environmental changes on phylogeographic patterns.

Complex impacts of climatic change are particularly exacerbated in freshwater ecosystems, as increasing temperature and aridity alters the stability and structure of hydrological systems (Middelkoop et al. 2001, Nijssen et al. 2001, Pinceel et al. 2013, Blöschl et al. 2019). With limited dispersal capability and reliance on available freshwater for survival, aquatic-dependent species demonstrate evolutionary associations with hydrological changes. Even minor alterations to hydrologic structure can have profound impacts on the evolution of a diverse array of freshwater taxa (Inoue et al. 2014, Thomaz et al. 2017, Wallis et al. 2017). For example, tectonic activity can reshape waterways, leading to river capture across new areas and shifting distributional patterns of water-dependent species (Murphy and Austin 2004). Thus, freshwater biodiversity functions as an important indicator of the impact of historical environmental changes.

An ideal system for studying biogeographic changes in southeast Australia is the southern pygmy perch, *Nannoperca australis* (Percichthyidae). This small-bodied (<80mm), habitat-specialist fish prefers slow flowing and vegetated ephemeral streams (Wedderburn et al. 2012, Hammer et al. 2013). It is distributed throughout the temperate southeast Australia region, occupying the MDB, coastal Victoria and northern Tasmanian rivers. Previous phylogenetic work indicated that southern pygmy perch from eastern Victoria, Flinders Island and north-eastern Tasmania belong to a genetically distinct cryptic species referred to as *Nannoperca* ‘flindersi’ (Unmack et al. 2013, Buckley et al. 2018). Estimates of divergence time using a biogeographic calibration point suggest this split occurred ∼6.1 Ma (Buckley et al. 2018), but the biogeographic forces driving this speciation remain unknown. Being an ancient lineage, *N. australis* has likely responded to a variety of environmental changes across inland and coastal habitats since the Miocene. Furthermore, *N. australis* is threatened, particularly within the MDB, due to extreme pressure from anthropogenic changes to water flow, introduced predators and contemporary climate change (Balcombe et al. 2011, Brauer et al. 2016). The low dispersal capability, effective population size and high genetic structure of the species makes their survivability of great concern (Brauer et al. 2016, Brauer et al. 2017). Here we used genome-wide data assess the relative roles of hydrological and coastal topographic changes as drivers of evolutionary diversification and lineage persistence. We hypothesised that demographic changes and lineage diversification linked to aridification would be older (Miocene – Pliocene) and stronger for populations from inland basins, whereas changes linked to eustatic variation would be comparatively younger (Pleistocene) and common for populations from coastal or island habitats. We tested the impact of these factors using a hierarchical framework that incorporated complex, hypothesis-driven coalescent modelling, model-free demographic analyses and spatial (species distribution) modelling.

## MATERIALS & METHODS

### Sample Collection and Genomic Library Preparation

A total of 109 samples across 21 known populations of *N. australis* and three populations *N.* ‘flindersi’ (*n* = 4 – 5 individuals per population) were collected. This sample spans the full geographic range of the species and includes at least one population from each management unit (MU) identified in previous genetic and genomic studies (Table 1; Unmack et al. 2013, Cole et al. 2016). The sister species *N. obscura* (Buckley et al. 2018) was included as the outgroup for phylogenetic analyses (*n* = 5). Specimens were collected using a combination of electrofishing, dip-, fyke- or seine-netting. Specimens (either caudal fin or entire specimen) were stored dry at −80°C at the South Australian Museum, or in 99% ethanol at Flinders University.

**Table 1:** Locality data for samples used in this study. Abbreviations described in the table were those used for further analyses, while *n* refers to the number of individuals sequenced per population. *N. obscura* samples were only included as an outgroup in the phylogenetic analysis.

| Species | Population | Abbreviation | Field code | *n* |
| --- | --- | --- | --- | --- |
| *N. australis* | Angas R., Strathalbyn | NauANG | F-FISH84 | 5 |
|  | Lake Alexandrina | NauALE | SPPBrA* | 4 |
|  | Middle Ck, Warrenmang, Avoca | NauAVO | F-FISH75: PU99-33SPP | 5 |
|  | Jew Harp Ck, Sidonia | NauJHA | F-FISH78: PU00-01SPP | 3 |
|  | Tributary to Seven Creeks | NauSEV | PU13-65SPP | 4 |
|  | Merton Ck, Goulburn Rvr. | NauMER | F-FISHY6: PU09-01SPP | 5 |
|  | Broken R., Lima South | NauBRO | F-FISHY6: PU09-02SPP | 5 |
|  | King R., Cheshunt, Ovens Rvr. | NauKIN | F-FISHY6: PU09-06SPP | 4 |
|  | Spring Ck, Mitta Mitta | NauSPR | F-FISHY6: PU09-13SPP | 5 |
|  | Gap Ck, Kergunyah, Kiewa | NauGAP | F-FISHY6: PU09-12SPP<br>F-FISH77: PU99-81SPP | 5 |
|  | Murray R. lagoon, Albury | NauALB | F-FISH53: IW94-47 | 4 |
|  | Coppabella Ck, Coppabella | NauCOP | F-FISH75: PU99-82SPP | 5 |
|  | Blakney Ck, Lachlan Rvr. | NauLRT | F-FISH98: LPP-* | 5 |
|  | Glenelg R., Glenisla | NauGRG | F-FISH78: PU0014-SPP | 5 |
|  | Merri R., Grassmere | NauMRG | F-FISH78: PU00-22SPP | 5 |
|  | Curdies R., Curdie | NauCRC | F-FISH78: PU00-24SPP | 4 |
|  | Gellibrand R. floodplain | NauGRF | F-FISH97: PU02-92SPP | 5 |
|  | Barongarook Ck, Colac | NauBAR | SPP08-13 | 4 |
|  | Mundy Gully | NauMG | F-FISHY8: PU08-11SPP | 4 |
|  | Gnarkeet Ck, Hamilton | NauGCH | F-FISHY2: PU00-27SPP | 4 |
|  | Wilsons Promontory | NauWP | F-FISH97: PU02-70SPP | 5 |
| *N. 'flindersi'* | Snowy R. lagoon, Orbost | NflSRLO | F-FISH77: PU99-85SPP | 5 |
|  | Flinders Island | NflFI | F-FISH84: FI-* | 4 |
|  | Anson R. tributary | NflANS | F-FISH82: HT-2* | 5 |
| *N. obscura* | Lake Alexandrina | Outgroup | YPBR* | 5 |
| **Total** |  | **24** |  | **109 (114)** |

DNA was extracted from muscle tissue or fin clips using a modified salting-out method (Sunnucks and Hales 1996) or a Qiagen DNeasy kit (Qiagen Inc., Valencia, CA, USA). Genomic DNA quality was assessed using a spectrophotometer (NanoDrop, Thermo Scientific), 2% agarose gels, and a fluorometer (Qubit, Life Technologies). All ddRAD genomic libraries were prepared in-house following Peterson et al. (2012), with modifications as described in Brauer et al. (2016).

Of the 109 samples, 73 were previously paired-end sequenced as part of a landscape genomics study (Brauer et al. 2016) using an Illumina HiSeq 2000 at Genome Quebec (Montreal, Canada). The additional 36 samples were single-end sequenced on an Illumina HiSeq 2500 at the South Australia Health and Medical Research Institute (SAHMRI).

### Bioinformatics

The resultant reads (forward reads only for paired-end samples) were filtered and demultiplexed using the ‘process_radtags’ module of Stacks 1.29 (Catchen et al. 2013), allowing ≤ 2 mismatches in the barcodes. Barcodes were removed and reads trimmed to 80□bp to remove low-quality bases from the ends. Cut reads were then aligned using PyRAD 3.0.6 (Eaton 2014), and further filtered by removing reads that had > 5□bp with a Phred score of < 20. Loci were retained at a minimum sequencing depth of 5 and occurring in at least ∼90% of samples (103). The final concatenated alignment contained 7,958 ddRAD loci and 45,104 SNPs.

### Phylogenetic Analysis

To determine evolutionary relationships as a basis for phylogeographic modelling, a maximum likelihood (ML) phylogeny was estimated using RAxML 8.2.11 (Stamatakis 2014) and the 7,958 concatenated ddRAD loci dataset. This was done using rapid hill-climbing and 1,000 resampling estimated log-likelihood (Pante et al. 2015) bootstraps under a GTR+ □substitution model. The resultant phylogenetic tree was visualised using MEGA 7 (Kumar et al. 2016) and rooted with *N. obscura* as the outgroup.

To determine if dendritic river hierarchy alone could explain phylogenetic patterns across the Murray-Darling Basin lineage, and to clarify coalescent models (see Results), linear correlations between genetic and riverine distance were estimated using StreamTree (Kalinowski et al. 2008). StreamTree models genetic divergence across a dendritic river system and assigns a cost to each riverine segment, comparing this modelled distance with the empirical data. While StreamTree is often used with pairwise F_ST_ values (e.g. Brauer et al. 2018) to assess contemporary spatial patterns, we opted to use uncorrected genetic distances (*p*-distance) as this more likely contains signal of historic patterns of divergence (Nei 2001). Pairwise *p*-distances between individuals were estimated using PAUP* 4 (Swofford 2002) and averaged per population using *R* for all 13 MDB populations.

### Divergence Time Estimation

We estimated divergence times using r8s 1.81 (Sanderson 2003). Given the lack of suitable fossils for pygmy perches (Buckley et al. 2018), we calibrated the node between *N. australis* and *N.* ‘flindersi’ at 5.9 – 6.1 Ma based on a previous estimate that includes all pygmy perch species (Buckley et al. 2018). Divergence times for each node were estimated using a penalized-likelihood model under a truncated Newton algorithm (Nash 2000), which uses a parametric branch substitution rate model with a nonparametric roughness penalty (Sanderson 2003). Cross-validation was used to determine the best value of the smoothing parameter for the roughness penalty between log_10_ 0 and log_10_ 100. The optimum smoothing parameter of log_10_ 41.00, with a chi-square error of −12836.285, was used to estimate divergence times between populations and higher order clades across the lineage.

### Ancestral Range Estimation

We used a phylogenetic tree-based method to estimate ancestral ranges across the maximum likelihood tree with the R package BioGeoBEARS (Matzke 2013). The maximum likelihood tree was collapsed down to individual populations using the R package ape (Paradis et al. 2004). Given the paraphyletic nature of the Albury population (NauALB) within the phylogenetic tree (Fig. 2), this population was pruned from the tree. The tree was then converted to ultrametric format using the divergence time estimates from r8s.

**Figure 2:**
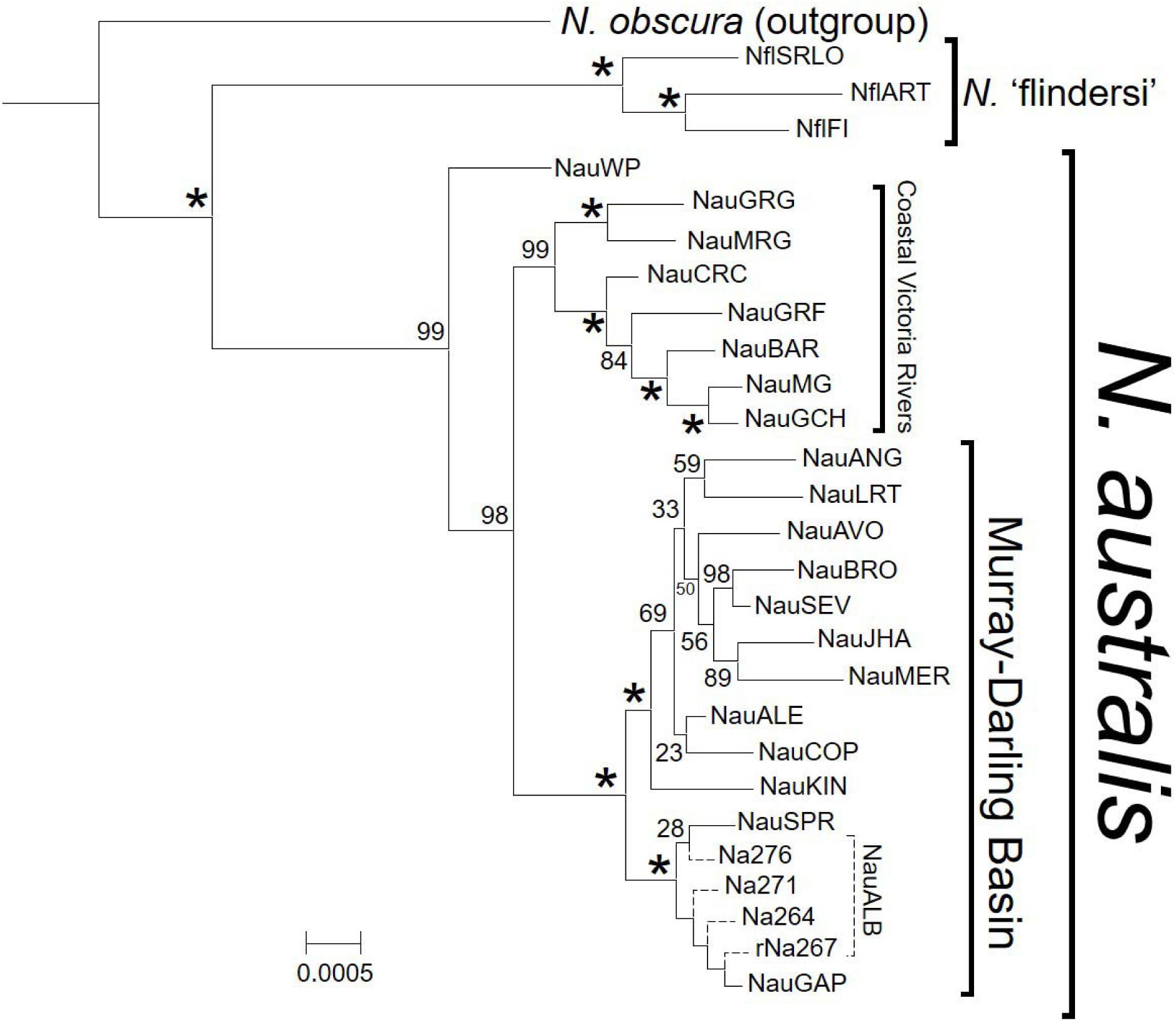
Maximum likelihood phylogeny of *N. australis* and *N.* ‘flindersi’ using 7,958 concatenated ddRAD loci containing 45,104 SNPs. As all samples within a population formed monophyletic clades (excluding NauALB, shown in dashed lines), the phylogeny was collapsed to individual populations. Nodes with 100% bootstrap support are indicated by asterisks. The tree was rooted using *N. obscura* as the outgroup. The full phylogenetic tree with all 119 samples is shown in Figure S1.

Tips were assigned to one of six main geographic regions based on hydrogeology (McLaren et al. 2011): the MDB, western Victoria coast (COAST), Wilson’s Promontory (WP), eastern Victoria (SRLO), Flinders Island (FLI) or Tasmania (ART). Individual *N.* ‘flindersi’ populations were assigned to unique geographic states given their current isolation and to allow for the explicit testing of vicariance vs. dispersal scenarios across the Bassian Isthmus. Ranges spanning multiple states were filtered to only those composed of neighbouring ranges (total number of possible ranges = 21). Furthermore, given the historical marine inundation of the MDB which would have precluded the presence of southern pygmy perch, time-stratification was used to exclude ‘MDB’ as a geographic state prior to 5 Ma or 3 Ma. These times reflect a conservative estimate of marine inundation (the most recent time at which marine sediments have been accurately identified within paleolake Bungunnia; McLaren et al. 2011) and a more relaxed estimate that is possibly the most recent time marine water could have been present within the basin. Ancestral ranges were estimated under all six available models (DEC, DIVA-LIKE and BAYAREA-LIKE, with and without founder-event speciation, +J). All models were run under both time-stratification scenarios and compared using the Akaike Information Criterion (AIC) within each set.

### Biogeographic Hypothesis Testing Using Coalescent Modelling

Specific hypotheses based on biogeographic events were tested using a coalescent framework within FastSimCoal 2.6 (Excoffier et al. 2013). These hypotheses expand on the broad interpretations based on phylogenetic analyses. Predicted changes in population history were based on the expected impact of biogeographic patterns previously identified within the literature (Table S2) and information from both prior and current phylogenetic and coalescent analyses (Buckley et al. 2018). These models were focused around particular divergence events across the lineage, with hypotheses built around the separation of major clades within the phylogeny (Fig. 2). Hypothesised biogeographical drivers of divergences included marine inundation, tectonic shifts and hydrological rearrangements. Specific biogeographic hypotheses for each divergence event, and their predictive impacts on the evolution and demography of southern pygmy perch, are described in Table S2 and Fig. S5.

### Estimating Effective Population Size Changes

As a model-free and exploratory approach to clarify demographic history, changes in effective population size (*Ne*) over time were estimated using the site-frequency spectrum (SFS) and coalescent modelling in a stairway plot (Liu and Fu 2015). Given the strong population genomic structure reported for southern pygmy perch (e.g. F_ST_ = 0 – 0.798; Brauer et al. 2016) including for populations sampled in this study, and to account for the biasing effect of population structure on the SFS (Stadler et al. 2009, Xue and Hickerson 2015), loci were re-aligned for each population independent of the rest of the dataset. As missing data can bias the distribution of the SFS (Shafer et al. 2017), only loci present in all samples for each population were retained. Unlinked biallelic SNPs from each independent alignment were then used to generate the single-population SFS (Fig. S4) using the same in-house script as above. Stairway plots were estimated assuming a mutation rate of 10^−8^ – 10^−9^ per site per generation (Stobie et al. 2018) and a generation time of one year (Humphries 1995).

### Species and Lineage Distribution Modelling

The distribution of the species was modelled using MaxEnt 3.4 (Phillips et al. 2017) and 19 BioClim variables from WorldClim v1.4 (Hijmans et al. 2005), summarising precipitation and temperature – two groups of climatic variables known to impact on local adaptation and distribution of southern pygmy perches (Brauer et al. 2016). To account for non-climatic environmental aspects that may limit the distribution of the species (Paz et al. 2015), elevation (extracted from the Etopo1 combined bathymetry and topography dataset; Amante and Eakins 2009) and topographic wetness index (twi, extracted from the ENVIREM database; Title and Bemmels 2018) were also included. Species occurrence data was collected from the Atlas of Living Australia (http://www.ala.org.au), with filtering for geographic accuracy and removing outliers based on known distributional limits (final data included 6,106 occurrences). However, this dataset did not include some localities within the Murrumbidgee River and mid-Murray River from which *N. australis* has been extirpated due to post-European settlement habitat modification across the MDB (Cole et al. 2016). Duplicates from the same coordinate point were removed to minimise the biasing effect of uneven sampling effort (Elith et al. 2011), reducing the dataset to 2,528 unique occurrences. Similar tests of spatial autocorrelation were performed for the 19 BioClim variables using a Pearson’s pairwise correlation test in SDMToolbox (Table S4; Brown et al. 2017). Highly correlated (*|r|* > 0.8) variables were removed to avoid overfitting of the model (Dormann *et al*. 2013), reducing the environmental data down to 9 bioclimatic variables and the two topographic variables (Table S5). A subset of 25% of occurrence sites were used to train the model.

Climatic data from the Last Glacial Maxima (LGM; 22 Ka) were extrapolated from the WorldClim 1.4 database (Hijmans et al. 2005) to project the historic distribution of the species. To evaluate environmental conditions more reflective of the divergence between the two species, the SDM was also projected back to the mid-Pliocene (∼3.2 Ma) using a subset of 6 of the previous 9 BioClim variables (excluding variables bio2, bio3 and bio6) from the PaleoClim database (Brown et al. 2018). The fit of each SDM was determined using the area under the receiver operating curve (AUC).

A lineage-specific distribution model (LDM) method described in Rosauer et al. (2015) was used to determine the relative distributions of each lineage over time; this was done with two ‘intraspecific’ lineages of *N. australis* and *N.* ‘flindersi’. A total of 72 site localities (*n* = 61 for *N. australis*; *n* = 11 for *N.* ‘flindersi’) were used based on genetic assignment to a ‘species’ within this study, as well as based on mitochondrial DNA results (Unmack et al. 2013). We estimated the LDMs for both species across all three time periods (current, LGM, and Pliocene). Although the location of intraspecific lineages is unlikely to remain constant in time, this method allows the inference of probable relative distributions of each ‘species’ under past climatic conditions.

## RESULTS

### Bioinformatics

A total of 340,950,849 raw sequenced reads were obtained from the 2 lanes of Illumina sequencing. Quality control and alignment resulted in a concatenated sequence dataset of 7,958 ddRAD loci with 45,104 variable sites (SNPs) and 30,485 parsimony-informative sites. This alignment had an average of 2.34% (±3.31% SD) missing data per individual. For coalescent analyses within FastSimCoal2, the outgroup was removed and loci realigned and SNPs re-called, generating a dataset of 8,022 ddRAD loci containing 38,287 SNPs and 22,820 parsimony-informative sites. SNPs from this dataset were reduced to a single SNP per ddRAD locus for estimating joint site frequency spectra, resulting in 7,780 biallelic SNPs.

### Phylogenetic Analysis

The maximum likelihood phylogeny (Fig. 2) separated *N.* ‘flindersi’ from the rest of *N. australis*, corroborating previous phylogenetic results (Unmack et al. 2013, Buckley et al. 2018). Within *N. australis*, three major lineages were delineated; one of the Wilson’s Promontory population (NauWP), one of coastal Victoria populations and another of populations within the MDB. The coastal Victorian lineage showed relatively stronger phylogenetic structure than the MDB lineage, with its easternmost populations diverging more recently compared to westernmost coastal Victorian populations (Fig. 4). The MDB clade, however, generally featured shorter branches and lower bootstrap support. Despite being geographically apart, lower MDB populations (Lake Alexandrina and Angas) shared a MRCA with upper Murray populations (Lachlan River and Coppabella Creek, respectively). The most basal clade of the MDB lineage contained populations from the upper Murray River (Spring Creek, Gap Creek, and Albury). Within this group, the Albury population was paraphyletic with the other two; this is expected based on previously described levels of admixture across the populations (Brauer et al. 2016). The breadth of the phylogenetic tree was well supported, with the majority of population-level and above divergences with bootstrap support of >80%.

The StreamTree results did not suggest that contemporary riverine hierarchy alone could explain patterns of historical phylogeographic divergence across the MDB (Fig. 3; R^2^ = 0.464). Assessing the fit of the StreamTree model by comparing the observed and expected genetic distance for each population individually demonstrated that this was likely driven by several outlier populations (NauANG, NauALE, NauALB and NauCOP) with significantly higher modelled genetic distance (Fig. S2). Removal of these four populations from the StreamTree model led to much higher correlation (R^2^ = 0.982) with similar genetic distance penalties for river segments common to both sets of populations (Fig. S3).

**Figure 3:**
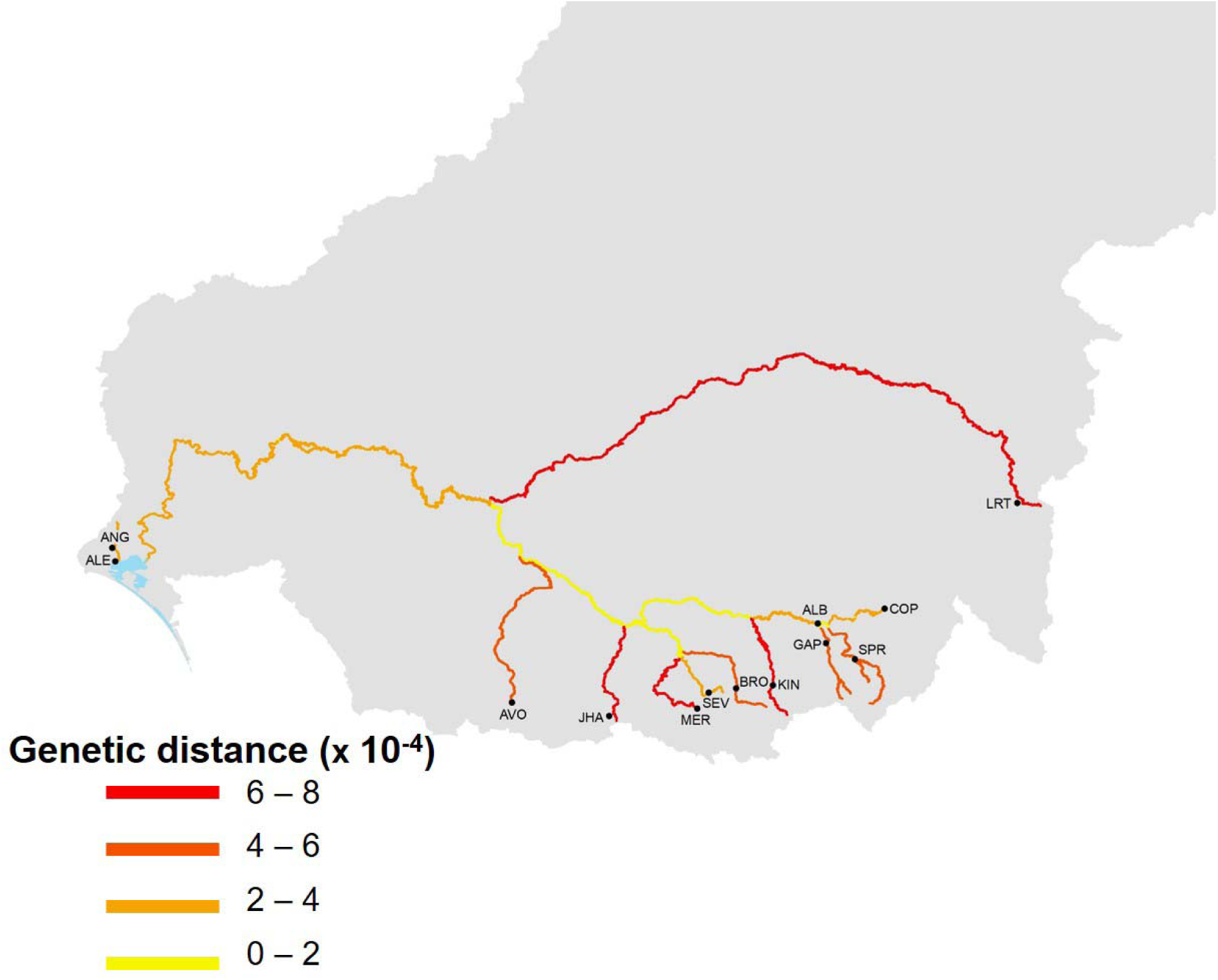
Dendritic riverine network of the Murray-Darling Basin (MDB), with streams colour-coded according to the StreamTree model that determines the contribution (as a penalty) of each segment in driving genetic divergence across the basin. Segments coloured in yellow confer little penalty (i.e. genetic divergence between populations at either end of the segment is low) whereas red segments confer higher genetic differentiation.

**Figure 4:**
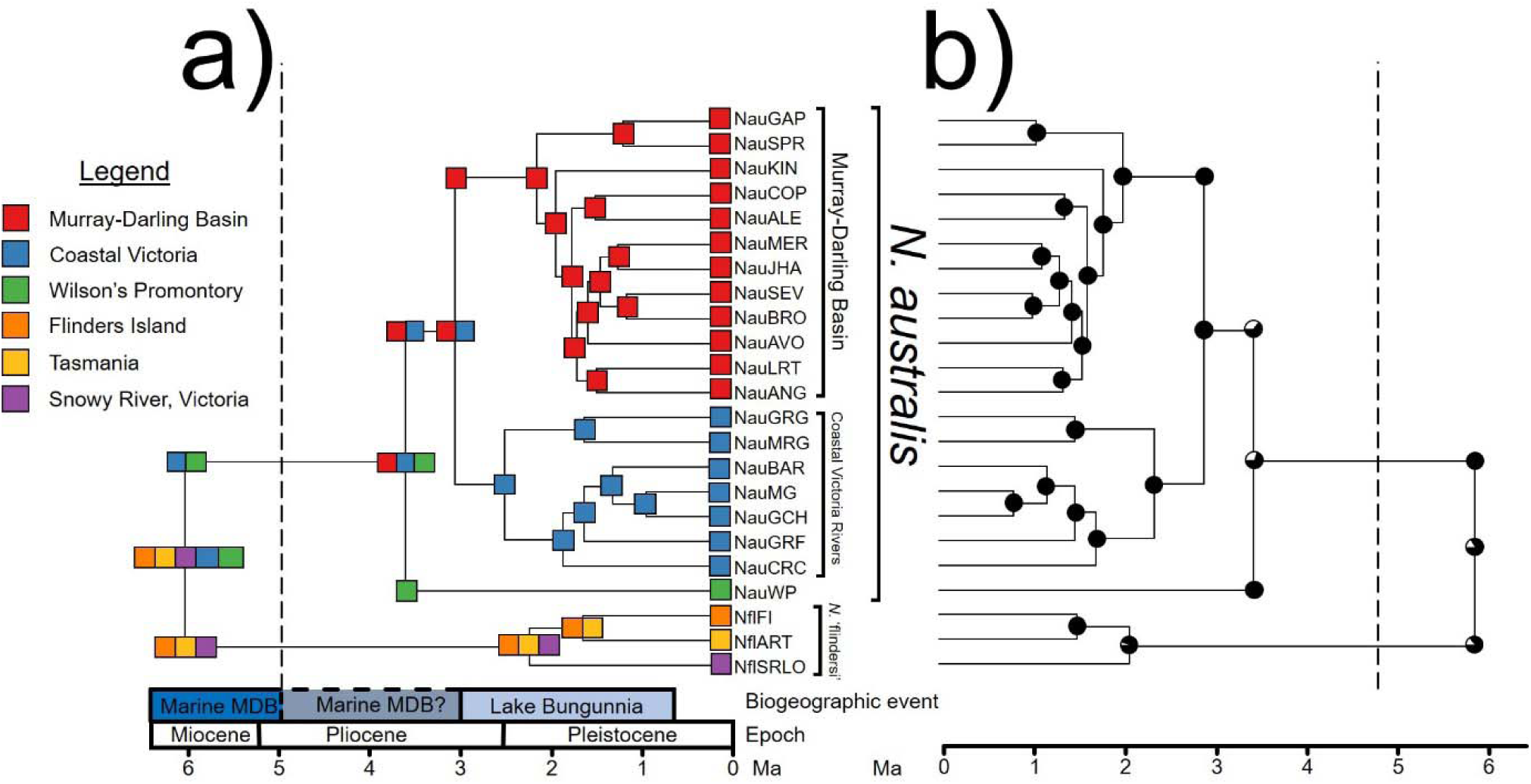
Most likely ancestral areas under the best supported model (DIVA-LIKE), with presence in the Murray-Darling Basin (MDB) excluded until 5 Ma (indicated by the dashed line). A biogeographic timeline of major alterations to the MDB is included for reference. Colours denote one of six contemporary areas, or ranges combining more than one area, as described by the legend. **a)** Most likely state at each node or branch where state changes occurred. **b)** Probability of the most likely state (black) for each node.

### Ancestral Range Estimation

Comparison of ancestral range estimations from BioGeoBEARS identified the DIVA-LIKE model as the best supported under both time-stratification scenarios (AIC = 24.16 and 59.1 for models excluding presence in the MDB until 5 Ma and 2 Ma, respectively). This model represents a likelihood approximation of the model implemented in DIVA (Ronquist 1997) which broadly considers the relative role of dispersal and vicariance (but not sympatric mechanisms) in driving biogeographic patterns (Matzke 2013). Although patterns were similar across both time-stratification criteria (Table S1), we choose to focus on the more conservative (5 Ma constraint) results given the lack of precision in determining the end of marine inundation into the MDB (McLaren *et al*. 2011). This DIVA-LIKE model demonstrated strong patterns of vicariance, with weak contributions of dispersal (*d* = 1.46 × 10^−2^) and extinction (*e* = 1 × 10^−12^) and all major geographic changes associated with vicariance events (Fig. 4). Including a parameter for founder event (+J) estimated very weak contributions of founding events and contributed to negligible change in log likelihood across either time-stratification scenario (Table S1). Ancestral states for major nodes were well resolved across the phylogeny.

### Biogeographic Hypothesis Testing

Comparison of biogeographic hypotheses under coalescent models clearly supported one model over others for each focal divergence event (Fig. 5): these results are detailed in Table S4. In general, most models including post-isolation gene flow were better supported than those without, and models based on vicariance due to hydrological changes were better supported than those invoking tectonic or dispersal mechanisms.

**Figure 5:**
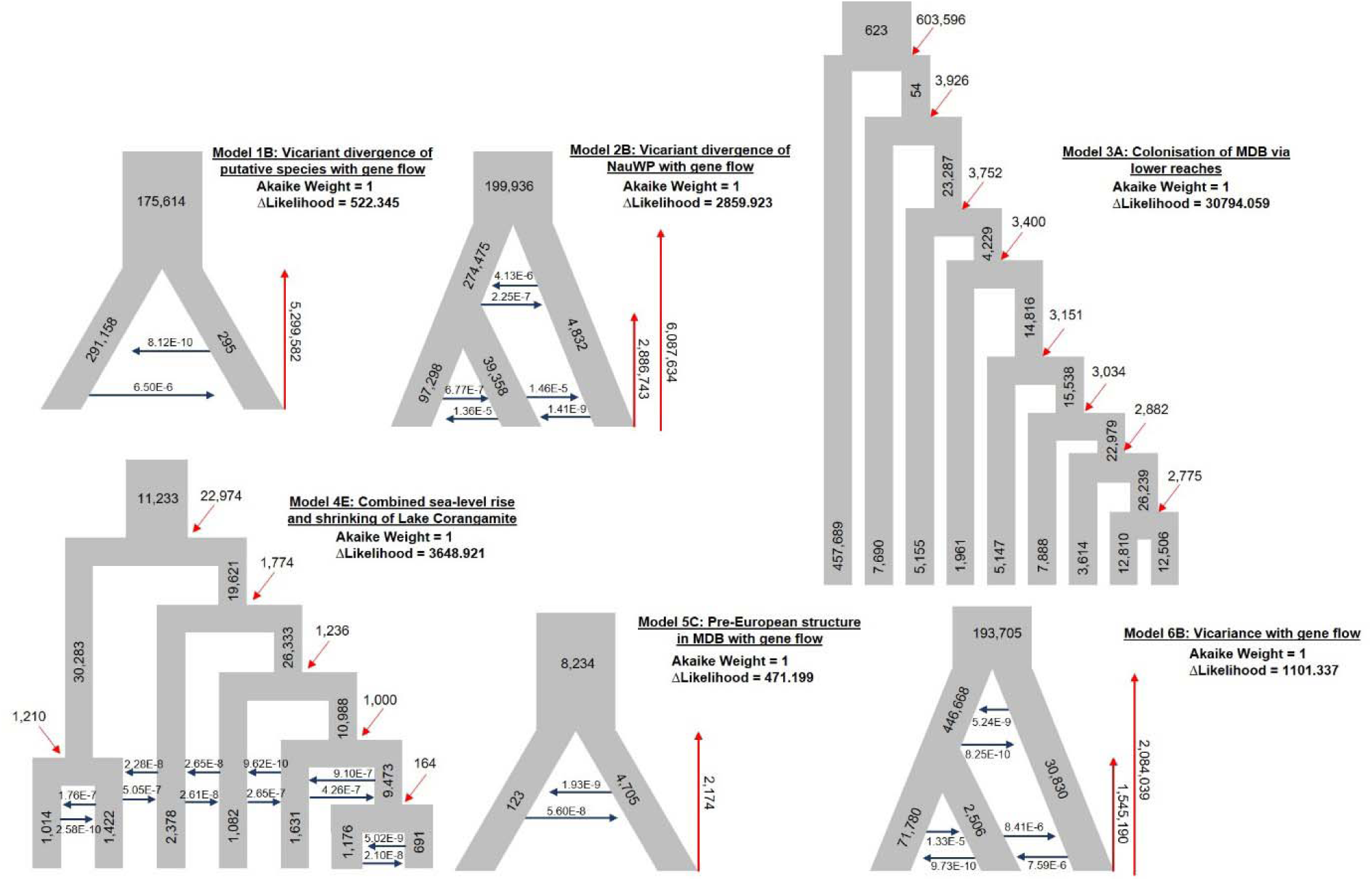
Representative diagrams of the best supported coalescent models under each model set. The full set of tested models and the biogeographic hypotheses underpinning them are described within the Supplementary Material. Red arrows denote divergence time parameters whilst blue arrows denote migration rate parameters. Population sizes are reported as the number of diploid individuals (N/2). Gene flow parameters are reported in terms of proportion of alleles moving in the direction of the arrow forward in time. ΔLikelihoods = difference between estimated (simulated) model likelihoods and observed (empirical) likelihoods.

#### Divergence of species

Coalescent models including gene flow between the two species after divergence was better supported than models without, suggesting that secondary contact occurred at some point after their initial divergence (Fig. 5; Model 1b). Models including bottlenecks did not have significantly higher support than those without, and inferred bottlenecks were small in magnitude. The initial divergence event between the two species was estimated at ∼6.2 Ma. These results suggest that range expansion and vicariant separation during the late Miocene drove speciation of *N.* ‘flindersi’.

#### Divergence of Wilson’s Promontory population

Coalescent models of the Wilson’s Promontory population and the two species separately also suggested likely gene flow across adjacent lineages, with divergence of the Wilson’s Promontory population occurring ∼3 Ma (Fig. 5; Model 2b). However, this gene flow was not indicative of a hybridisation event and models simulating the WP population as resulting from gene flow from either population coalesced nearly all alleles into the *N. australis* ancestral population. Thus, NauWP represents an anciently diverged population of *N. australis* that was isolated through vicariance, possibly due to either marine inundation of the peninsula or tectonic changes across the region.

#### Divergence of major N. australis lineages

Testing migration pathways from the ancestral coastal population into the MDB suggested that colonisation most likely occurred through the lower sections of the MDB and upstream into the upper reaches (Fig 5.; Model 3a). The timing of this migration event would pre-date ∼600 Ka, the estimated time of divergence between the coastal and MDB populations within the best supported model. However, this model had only marginally better likelihood than one estimating the divergence time of the coast and MDB populations at ∼1.2 Ma, suggesting that this estimate might not be overly precise. Regardless, the biogeographic models suggest that migration facilitated by the presence of paleo megalake Bungunnia allowed the ancestral southern pygmy perch to enter the MDB following the withdrawal of marine water from the basin.

#### Divergence within coastal Victoria lineage

The coalescent model accounting for the effects of both sea-level changes leading to isolation of rivers and the shrinking of Lake Corangamite isolating eastern lineages was better supported than models without these factors, or models only considering one (Fig. 5; Model 4e). This suggests that while sea-level changes likely isolated many of the populations from one another and prevented dispersal across river systems, the expanded Lake Corangamite continued to facilitate gene flow across some of the more eastern lineages.

#### Divergence within MDB lineage

Coalescent modelling of the MDB lineage suggested that some phylogeographic structure pre-dated European settlement, with the divergence of the most basal lineage (containing the Spring Creek, Gap Creek and Albury populations) estimated to have occurred ∼2 Ka, albeit with low levels of gene flow since divergence (Fig. 5; Model 5c). Models partitioning putatively upper and lower populations into single demes did not converge, likely reflecting their paraphyletic nature.

#### Divergence within N. ‘flindersi’

Coalescent models including migration between adjacent populations gave much greater likelihood estimations than models without gene flow, suggesting that migration between lineages had occurred in the past (Fig. 5; Model 6b). Including bottlenecks indicative of a dispersal event did not improve likelihoods, supporting a range expansion and vicariance scenario. The central population of Flinders Island had a much smaller population size than either of the other two populations. Divergence time estimates for between populations suggest a relatively ancient split, with the Snowy River population separating from the other two lineages ∼1.5 Ma and the secondary split between Flinders Island and Tasmania at ∼1.3 Ma.

#### Historical Demography Reconstruction

One-dimensional site-frequency spectra estimated from single-population SNPs used a mean of 3,527 (1,045 – 7,389) SNPs (Fig. S4). Stairway plots indicated significant declines in many populations of southern pygmy perch over the last 1 Myr (Fig. 6). For many of these populations, gradual and concordant bottlenecks were apparent within clades. Non-declining populations were typically relatively stable, with none demonstrating an increase in *Ne* over this time. Almost all populations across both putative species demonstrated population growth deep in the past (∼ 1 Ma), although this may reflect fewer ancient coalescent events within the data.

**Figure 6:**
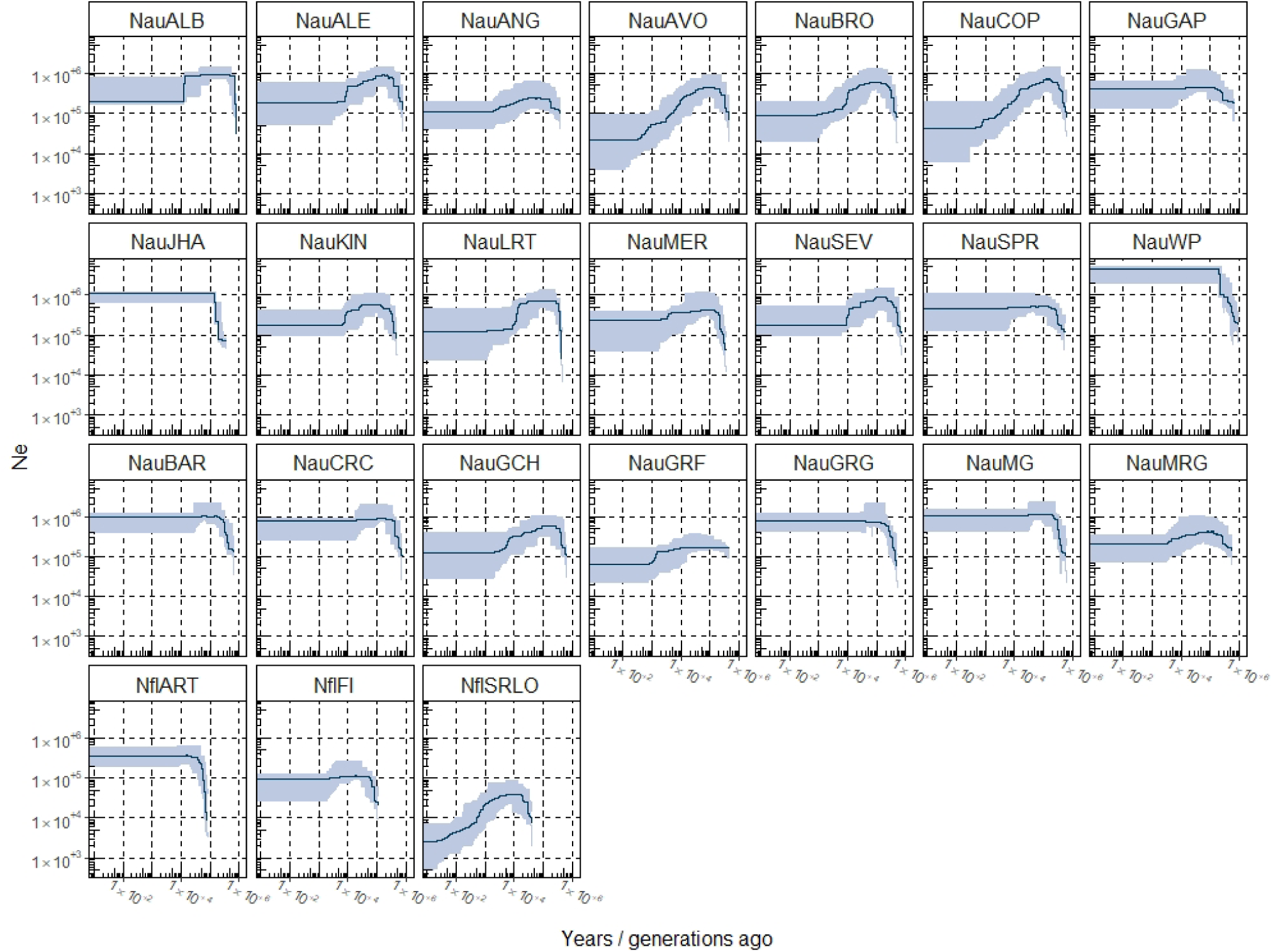
Stairway plot reconstructions of demographic history for individual populations. Both axes are reported in log_10_ scale. Dark blue lines indicate medians with 95% confidence intervals shaded. Top two rows = Murray-Darling Basin (MDB) and WP populations; third row = coastal Victoria populations; bottom row = *N.* ‘flindersi’ populations.

Within *N.* ‘flindersi’, the mainland population of NflSRLO demonstrated a strong decline in *Ne* starting at ∼10 Ka which contrasted with the more stable demographic histories of the island populations. This decline resulted in much lower estimates of recent *Ne* than the island counterparts, with the Tasmanian population NflART showing the highest consistent *Ne* of all *N.* ‘flindersi’ populations.

Demographic histories were variable across populations of *N. australis*. Within the coastal ESU, most populations demonstrated relatively stable demographic histories, with weak declines in *Ne* originating at ∼100 Ka in NauMRG and NauGCH, and at ∼10 Ka in NauGRF. This contrasted with populations across the MDB, where significant population declines at ∼100 Ka were observed in the majority of populations. Nevertheless, a few populations also demonstrated stable or weakly declining *Ne* over time, with NauJHA and outlier showing significant growth at ∼ 100 Ka followed by stable *Ne.* However, this is likely driven by the lower sample size (*n* = 3) within this population compared to others across the MDB, resulting in few inferred coalescent events across the tree and a biased SFS. The highly divergent Wilson’s Promontory population showed a sharp population increase at ∼100 Ka followed by relatively stable and high *Ne* over time. Although all population stairway plots inferred no changes in *Ne* <5 Ka, this likely reflects a lack of very recent coalescent events within each population owing to small sample size.

#### Species and Lineage Distribution Modelling

Species distribution modelling for southern pygmy perch based on the nine uncorrelated BioClim and 2 topographic variables effectively predicted the contemporary distribution for the species, showing highest habitat suitability along the Victorian coast, southern MDB and in northern Tasmania (including King and Flinders Island; Fig. 7a). However, this SDM likely underestimates presence of *N. australis* within the connective center of the MDB, where downstream migration would have facilitated a mosaic of intermediate populations prior to extensive flow abstraction and regulation over the last 200 years (Cole et al. 2016).

**Figure 7:**
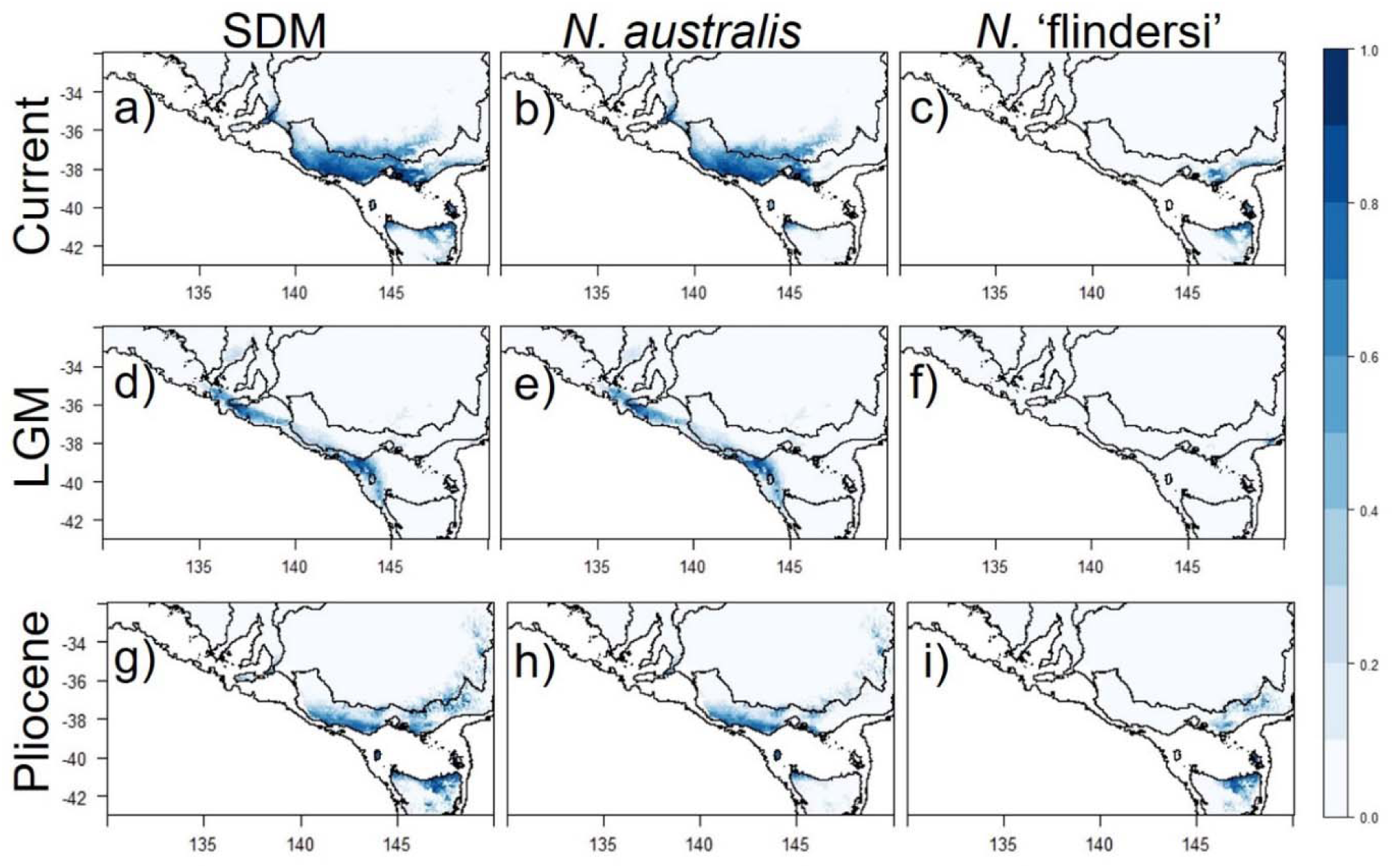
Species distribution models for all lineages and lineage-specific distribution models for each putative species based on 9 bioclimatic and 2 topographic variables. **a-c)** Distributions under contemporary climate conditions. **d-f)** Distributions under last glacial maximum (LGM; 22 Ka) climate. **g-i)** Distributions under mid Pliocene (3.2 Ma) climate conditions.

Historic projections of the SDM for both species together highlighted two glacial refugia, one along the western coast of the distribution and another small and isolated refugium closer to the southeast corner of the mainland. Together, these results indicate a significant expansion in suitable habitat following the LGM (Fig. 7d). The AUC of the model was estimated at 0.908, indicating a good fit of the model to the data.

Lineage distribution models under each time period demonstrated the disjunct spatial nature of the two species. Under contemporary conditions, the LDMs showed a geographic break near Wilson’s Promontory, albeit with overlap at intermediate localities (Fig. 7b-c). Additionally, the northern Tasmanian coastal habitat was delimited into two equally sized sections for each species. During the LGM, the disjunct LDMs of the two species in the east and west portion of the range supported their long-term isolation and provided evidence for an environmental barrier preventing the contact between the species (Fig. 7e-f). The Pliocene projection also showed refugial habitat along the Victorian coast and within the upper MDB (Fig. 7g). Similarly to contemporary conditions, the LDMs showed a narrow division between species in northern Tasmania and central Victoria (Fig. 7h-i). The subset data used for the Pliocene projection had marginally weaker support, with an AUC of 0.887.

## DISCUSSION

Establishing how aquatic-dependent lineages responded to past hydroclimatic changes contributes to our understanding about their contemporary ecological requirements and to predicting demographic responses under ongoing climate change. This study demonstrates the overarching impacts of varying hydrology due to Plio-Pleistocene climatic change (e.g. reduction of lake systems and rearrangement of river networks) on the evolution and diversification of a temperate freshwater-dependent fish clade. However, coalescent analyses and species distribution modeling show that the evolutionary consequences of major shifts in sea level and hydroclimatic conditions varied substantially between coastal and inland environments. Aridification altered the demography of populations from inland river systems, whilst eustatic changes and marine inundation were major evolutionary drivers of populations from coastal freshwater landscapes. Our findings suggest that the long-term impact of key environmental changes associated with anthropogenic climate change, such as increases in aridification and sea level, might vary substantially for the same lineage, both spatially and across landscape types.

### Aridification Drives Phylogeographic Structure of Inland Basins

Aridification of the Australian continent has dramatically altered the identity and stability of ecosystems (Hawlitschek et al. 2012), particularly in the formation of the arid zone (Byrne et al. 2008) and reduction of temperate and wetter habitats (Crisp et al. 2004, Byrne et al. 2011). Particularly for freshwater species, increasing aridification since the Pliocene may be responsible for a number of divergent clades, with water availability and river networks being critical for the long-term survival and evolution of freshwater lineages (Faulks et al. 2010, Beheregaray et al. 2017).

Aridification and tectonics through the formation and demise of paleo-megalake Bungunnia was a major event impacting the evolutionary history of inland lineages of *N. australis*. Lake Bungunnia initially formed ∼3 Ma when tectonics shifts along the Padthaway High resulted in significant uplift across the region, damming the ancestral Murray River which approximately aligned with the current Glenelg River (McLaren et al. 2011, Waters et al. 2019). For many freshwater taxa across the southeast of the continent, isolation of lineages between the MDB and the southwest Victoria (SWV) drainages has been associated with this tectonic shift in the Pliocene (Murphy and Austin 2004, Waters et al. 2019). Similar interpretations of tectonic changes influencing river capture have been proposed for movement across other sections of the Great Dividing Range into the MDB (McGlashan 2001, Murphy and Austin 2004, Cook et al. 2006, Faulks et al. 2010). The isolating effect of the tectonic changes between the two drainages is likely reflected in the strong differentiation between clades and the ancient nature of the phylogenetic-based divergence time of 3.03 Ma.

Coalescent modelling instead points to more recent divergence (603 Ka), suggesting the possibility that the edges of Lake Bungunnia at its largest extent could have acted as suitable habitat for southern pygmy perch and potentially facilitated further dispersal into the inland basin. With the eventual demise of the lake ∼700 Ka (McLaren et al. 2011), this secondary bout of isolation disconnected the two basins fully, probably accounting for the results of coalescent models. Similar patterns of initial isolation by vicariance during the Pliocene, followed by Pleistocene secondary contact across the Great Dividing Range and into the MDB were observed within mountain galaxias (*Galaxiis oliros* and *Galaxiis olidus*), which share comparable ecological constraints to southern pygmy perch (Waters et al. 2019). Similar to Lake Bungunnia, the reduction of Lake Corangamite to one seventh of its original size over the course of the Holocene (White 2000) isolated several eastern coastal Victoria *N. australis* populations. The reduced size (∼160 km^2^) and hypersalinity (>50 g/L) of Lake Corangamite likely prevents connectivity between these populations under contemporary conditions (Williams 1995, White 2000).

Within the MDB clade there was weak evidence for historic phylogeographic structure, with coalescent models suggesting divergences dating as ∼2 Ka. Correlating contemporary river structure and genetic distance *per se* did not predict genetic divergence between populations across the MDB. A combination of extensive flow and habitat modification since European settlement and naturally complex metapopulation dynamics (Brauer et al. 2016) are probably better proxies for contemporary patterns observed within the MDB. This was reflected in the stairway plots, which showed a number of populations declining over the last 1 Myr but with variance in demographic histories across the basin. The most likely demographic scenarios include multiple waves of dispersal and colonisation, possibly in response to local extinction or during rare environmental events such as flooding, which altered patterns of genetic divergence.

### Eustatic Changes Drive Phylogeography and Speciation Along Coastal Habitats

Sea level changes associated with interglacial periods and more arid climates played a significant role in the divergence of coastal lineages. Marine inundation across the East Gippsland region during the Mio-Pliocene (∼6 Ma), prior to the climatic cycles of the Pleistocene, likely drove the initial divergence and speciation of *N.* ‘flindersi’. Marine sediments and low elevation of the region indicates that this marine inundation was significant (Gallagher et al. 2001, Holdgate et al. 2003), and was correlated with the onset of major aridification in the continent (Garrick et al. 2004, Faulks et al. 2010, McLaren and Wallace 2010). Ancient marine inundation of East Gippsland has been proposed to influence vicariant speciation in various terrestrial species (Chapple et al. 2005, Norgate et al. 2009). This low-lying region approximately forms the interface between the distribution of the two putative study species (Fig. 1b) and the timing of this inundation corresponds well with the estimated molecular clock-based divergence time. This period of sea level rise is also associated with inundation of the lower parts of the MDB which ostensibly precluded the presence of pygmy perch (McLaren et al. 2011).

Previous hypotheses of the mechanisms driving the initial divergence of *N.* ‘flindersi’ have suggested that the separation of drainages by tectonic shifts across the region (Dickinson et al. 2002, Gallagher et al. 2003) isolated populations following a dispersal event facilitated by river capture or flooding (Unmack et al. 2013). Regardless of the mechanism, divergence between Tasmanian and mainland lineages prior to the Pleistocene has been reported for birds (Lamb et al. 2019), lizards (Dubey and Shine 2010, Chapple et al. 2011, Kreger et al. 2019), butterflies (Norgate et al. 2009) and other freshwater fish (Coleman et al. 2010), suggesting that climatic oscillations during the LGM alone did not drive the speciation of *N.* ‘flindersi’.

More recent sea-level changes also likely impacted within-basin phylogeographic patterns. Within *N.* ‘flindersi’, relatively ancient estimates of divergence times between populations (1.5 – 2 Ma) suggested that early glacial cycles of the Plio-Pleistocene resulted in strong differentiation. However, coalescent models suggested that gene flow across these disparate populations was possible during glacial maxima. At lowered sea levels, river systems occupied by *N.* ‘flindersi’ all drained eastward towards the continental shelf (Unmack et al. 2013), with shorter overland distances between river mouths than today (Fig. 1b). Given the presence of a small glacial refugia in the far eastern extreme of the distribution, gene flow may have resulted from contraction into a singular locale followed by expansion back across the Bassian Isthmus during more favourable environmental conditions (Lambeck and Chappell 2001).

Within the coastal *N. australis* lineage, isolation of river catchments during aridification in the Pleistocene led to the strong structure observed within the phylogenetic tree. During glacial maxima, lowered sea levels significantly increased the extent of the mainland Australian coastline, particularly across the southeast corner (Williams et al. 2018). Although it does not appear that the current rivers of coastal Victoria ever fully connected together before meeting the shoreline (Unmack et al. 2013), climatic modelling has suggested that the low topographic relief and evaporation across this region would have allowed overland networks to form through small lakes and floodplains (Williams et al. 2018). Sea-level rise and aridification during the Pleistocene inundated much of this habitat and subsequently isolated river systems from one another. Thus, along coastal Victoria aridification and eustatic changes demonstrated interactive effects on phylogeographic patterns. The more historic nature of divergences within the coastal lineage compared to the MDB lineage were corroborated by the well-resolved phylogenetic structure and relative stability of population sizes over time across the clade.

### Identity and Maintenance of Cryptic Species, N. ‘flindersi’

Although the initial divergence between *N. australis* and *N.* ‘flindersi’ was associated with older biogeographic events during the Miocene, our results indicated weak post-divergence gene flow between the two species. Distribution modelling indicated a likely overlap in suitable habitat under contemporaneous conditions across the Victorian and Tasmanian coastlines, with little divergence in environmental ranges between the two species (Fig. S6). However, environmental changes during glacial maxima likely caused the two species to retract to isolated refugia. These factors together suggest a history of alternating periods of isolation and connectivity during glacial cycles, with isolated glacial refugia and weak interspecific interglacial gene flow limited to a narrow hybrid zone at the point of contact. Other studies of terrestrial species diversification across recurrently connected islands suggest patterns of gene flow in accordance with lower sea levels (Paulay and Meyer 2002, Jordan and Snell 2008, Parent et al. 2008, Papadopoulou and Knowles 2017). In this case, gene flow with *N. australis* does not appear to have impeded divergence and our results support the previous denotation of *N.* ‘flindersi’ as an independent species (Unmack et al. 2011, Unmack et al. 2013, Buckley et al. 2018).

### Implications for Conservation Management

Southern pygmy perch are currently listed as Near Threatened on the International Union for Conservation of Nature (IUCN, 2019) and Vulnerable or Endangered within state government management lists (Hammer et al. 2013). Ongoing conservation management has sought to recover their numbers, particularly across the MDB (Attard et al. 2016, Brauer et al. 2016, Cole et al. 2016). Across the species range, a number of clades have been identified and used as the basis for management practices (Unmack et al. 2013, Cole et al. 2016). Traditionally, conservation managers have adopted a ‘local is best’ paradigm, maintaining independence of populations within captive breeding and translocation programs to prevent outbreeding depression (Frankham et al. 2011, Love Stowell et al. 2017). However, the complex nature of the southern pygmy perch populations across the MDB indicates a history likely dictated by metapopulation dynamics with natural patterns of local extinction, recolonization and sporadic gene flow (Cole et al. 2016). A growing body of literature suggests that the propensity and magnitude of outbreeding depression has been overestimated (Frankham et al. 2011). Given the low levels of genetic diversity and high imperilment of MDB populations (Brauer et al. 2016), as well as the recent pattern of within-basin divergence detected here, we argue that *in situ* and *ex situ* conservation efforts should use a basin-wide context when selecting populations as source for demographic and genetic rescue (e.g. captive breeding and translocations).

### Implications Under Climate Change

Historic climatic fluctuations have often been used to predict future responses to anthropogenic climate change (MacDonald et al. 2008, Dawson et al. 2011). Primarily, these studies have focused on how species ranges and survival have responded to changes in temperature, precipitation and sea level. Correspondingly, however, many bioregions across the world are expected to increase in aridity with ongoing climate change (Christensen et al. 2007), impacting on the stability and structure of freshwater ecosystems globally (Middelkoop et al. 2001, Nijssen et al. 2001), including for the MDB (Cai and Cowan 2008, Pittock and Finlayson 2011). Additionally, drought events are projected to occur at higher frequency and with higher severity within Australia (Christensen et al. 2007). However, impacts of climatic change on the availability and reliability of water resources are uncertain (Middelkoop et al. 2001), as it is also the case on the influence of hydrological change on the evolution and persistence of species. Projections of sea level rise associated with melting glacial ice similarly predict major inundation of coastal habitats globally (Rotzoll and Fletcher 2012). This poses a threat to freshwater species not adapted to high salinity, and salinification of ecosystems to more estuarine or marine environments across coastal regions threatens swathes of biodiversity (Courchamp et al. 2014). Indeed, terrestrial species extinctions have already been directly linked to inundation of island habitats (Waller et al. 2017).

Our study highlights how spatial variation in the role and extent of environmental changes may result in variable impacts on the demography, distribution and divergence of populations. Particularly, we show how aridification of inland waterbodies and sea level rise leading to marine inundation of habitat impacted different regions across the distribution of a freshwater fish, operating on different timescales and to different extents. These environmental changes caused significant divergence across the clade, resulting in a hierarchy of lineages spanning from a cryptic species to intraspecific clades. While further increases in temperature will directly impact on the long-term survival of many species broadly, additional impacts on hydrological systems through aridification will have compounding effects on freshwater species. Our findings suggest that ongoing impacts from anthropogenic climate change may be complex in nature and vary across biogeographic regions depending on the role and identity of environmental forces that operate locally. We suggest that future management scenarios should consider this spatial variation in prediction of responses to climate change, particularly in how specific aspects (e.g. aridification, sea level changes) may act heterogeneously across species distributions.

## ACKNOWLEDGEMENTS

This work was supported by an Australian Research Council grant (FT130101068 to L.B.B.). We acknowledge researchers that provided samples or participated in field expeditions, especially Mark Adams. This work received logistic support from Flinders University, University of Canberra and the South Australian Museum.

## DATA ACCESSIBILITY

The sequence alignment used for phylogenetic-based analyses is available in NEXUS format in Dryad (**Dryad ref here**).

## AUTHOR CONTRIBUTIONS

S.B. contributed to all sections of data analysis as well as drafting the manuscript. L.B.B. designed the study, obtained resources, and helped with manuscript drafting. C.B. generated the data. P.U. and M.H. contributed with samples and field expertise. All authors contributed to the interpretation of results and critically revised the manuscript.

