## Supplementary Information for "The roles of aridification and sea level changes in the diversification and persistence of freshwater fish lineages"

Methods

*Biogeographic hypothesis testing framework using coalescent modelling*

We designed a set of coalescent models within FastSimCoal2.6 to test various hypotheses of biogeographic mechanisms and their impact on the evolution and divergence of southern pygmy perch populations. Sets of models were built around competing hypotheses that could explain major divergences across and within the major clades of the species pair (Fig. 1A). These hypotheses were based on information obtained from the topology of the phylogenetic tree (Fig. 2), estimates of divergence times (Fig. 4), and various studies on the environmental development of the region. Details of the biogeographic hypotheses underpinning the models are listed in Table S2. Given the high number of parameters required to estimate the number of populations and their associated demographic histories, some models involved amalgamating individual populations into a single coalescent deme. Although this potentially biases individual site frequency spectra, this reduction in parameterisation was required to feasibly run the models and all models across a given set featured the same population definitions. Thus, we do not expect that this reduction significantly influenced the *comparative* likelihoods of models across a set.

The joint site frequency spectrum (jSFS) was estimated by subsampling from the 7,780 shared SNPs to the populations specific to each model: as the total number and relevance of populations changed between different model hierarchies, this resulted in variables numbers of variable sites. All models were estimated under 40 ECM cycles with 500,000 simulations each and model fit was evaluated using AIC and Akaike Weights (Table S3), with the best supported model for each set chosen to represent the biogeographic event that best described the appropriate section of the tree.

Given the sheer number of models tested and their individual complexities, we did not compute confidence intervals through re-estimation per model. Instead, we focus on the comparative likelihoods of the demographic scenarios described by the models and treat individual parameter values with caution. Thus, parameters should be interpreted only in a relative context within each model or across sets of models.

**Table S1:** Comparison of ancestral range estimation model likelihoods across six biogeographic models and two time stratification scenarios, and including a constraint on maximum range size. The most supported models are indicated by asterisks.

| **Constraint** | **Model** | **Log likelihood** | **Dispersal (*d*)** | **Extincton (*e*)** | **Found event (*J*)** | **AIC** | **Relative AIC** |
| --- | --- | --- | --- | --- | --- | --- | --- |
| MDB excluded prior to 5 Mya | DEC | -17.6 | 0.030 | 1.0 x 10^-12^ | 0 | 39.19 | 0.0004 |
|  | DEC+J | -16.61 | 1.0 x 10^-12^ | 1.0 x 10^-12^ | 0.013 | 39.23 | 0.0004 |
|  | DIVALIKE | -10.08 | 0.015 | 1.0 x 10^-12^ | 0 | 24.16 | 0.73***** |
|  | DIVALIKE+J | -10.08 | 0.015 | 1.0 x 10^-12^ | 1.0 x 10^-5^ | 26.16 | 0.27 |
|  | BAYAREALIKE | -30.29 | 0.036 | 0.21 | 0 | 64.57 | 1.2 x 10^-9^ |
|  | BAYAREALIKE+J | -20.08 | 1.0 x 10^-12^ | 1.0 x 10^-12^ | 0.029 | 46.16 | 1.2 x 10^-5^ |
| MDB excluded prior to 5 Mya + max range = 2 areas | DEC | -23.07 | 0.059 | 1.0 x 10^-12^ | 0 | 50.14 | 0.0028 |
|  | DEC+J | -19.1 | 1.0 x 10^-12^ | 1.0 x 10^-12^ | 0.026 | 44.21 | 0.054 |
|  | DIVALIKE | -17.58 | 0.059 | 1.0 x 10^-12^ | 0 | 39.15 | 0.67***** |
|  | DIVALIKE+J | -17.55 | 0.039 | 1.0 x 10^-12^ | 0.008 | 41.1 | 0.25 |
|  | BAYAREALIKE | -32.1 | 0.093 | 0.13 | 0 | 68.2 | 3.30 x 10^-7^ |
|  | BAYAREALIKE+J | -20.08 | 1.0 x 10^-12^ | 1.0 x 10^-12^ | 0.029 | 46.16 | 0.02 |
| MDB excluded prior to 2 Mya | DEC | -30.35 | 0.050 | 0.062 | 0 | 64.69 | 0.042 |
|  | DEC+J | -30.35 | 0.050 | 0.062 | 1.0 x 10^-5^ | 66.69 | 0.016 |
|  | DIVALIKE | -27.55 | 0.040 | 0.041 | 0 | 59.1 | 0.69***** |
|  | DIVALIKE+J | -27.55 | 0.040 | 0.041 | 1.0 x 10^-5^ | 61.1 | 0.25 |
|  | BAYAREALIKE | -38.91 | 0.063 | 0.23 | 0 | 81.82 | 8.0 x 10^-6^ |
|  | BAYAREALIKE+J | -36.64 | 0.037 | 0.080 | 0.023 | 79.28 | 2.9 x 10^-5^ |

**Table S2:** Biogeographic hypothesis testing framework for coalescent modelling in FastSimCoal2

| **Model** |  | **Biogeographic mechanism** | **Reference(s)** | **Fixed/constrained parameters** | **Total number of parameters** |
| --- | --- | --- | --- | --- | --- |
| 1 | A | Range expansion and vicariant separation due to early cycles of marine inundation in E Gippsland | Chapple *et al.* 2005  Norgate *et al.* 2009  Coleman *et al.* 2010 | N/A | 4 |
|  | B | Range expansion and vicariant separation due to early cycles of marine inundation in E Gippsland with post-isolation migration | Chapple *et al.* 2005  Norgate *et al.* 2009  Coleman *et al.* 2010 | M_1_-M_2_ = 10^-10^ – 10^-5^ | 6 |
|  | C | Dispersal into Eastern Victoria after withdrawal of Eastern Gippsland inundation | N/A | Bottleneck | 6 |
|  | D | Post-inundation dispersal with post-isolation migration between species | N/A | Bottleneck  M_1_-M_2_ = 10^-10^ – 10^-5^ | 8 |
| 2 | A | Vicariant separation of NauWP due to tectonics, marine inundation or volcanism | Dickinson *et al.* 2002  Gallagher *et al.* 2001  Holdgate *et al.* 2003  Lesti *et al.* 2008 | T_WP_ = 4.5 – 2, 6 – 2, or <5.3 Mya, respectively. | 7 |
|  | B | Vicariant separation of NauWP with post-isolation migration | N/A | M_1_-M_6_ = 10^-10^ – 10^-5^ | 13 |
|  | C | NauWP is 50% hybrid lineage of *N. aus* and *N.* ‘fli’ | N/A | Coal_NauWP→Fli_ = 0.5; Coal_NauWP→Aus_ = 0.5 | 7 |
|  | D | NauWP is hybrid lineage of *N. aus* and *N.* ‘fli’ with variable contributions of gene flow between species | N/A | Coal_NauWP→Fli_ = 1^-10^ - 0.5 Coal_NauWP→Aus_ = 1^-10^ - 0.5 | 9 |
|  | E | NauWP is hybrid lineage of *N. aus* and *N.* ‘fli’ with variable contributions of gene flow between species with additional post-isolation migration | N/A | Coal_NauWP→Fli_ = 1^-10^ - 0.5 Coal_NauWP→Aus_ = 1^-10^ - 0.5  M_1_-M_6_ = 10^-10^ – 10^-5^ | 15 |
| 3 | A | Dispersal through the lower reaches of the MDB (via Lake Bungunnia) | McLaren *et al.* 2012 | T_COAST_ = 700 Kya, 1.2 Mya, 3 Mya. | 25 |
|  | B | Dispersal through the middle reaches of the MDB (across the GDR) | Unmack *et al.* 2001 |  | 25 |
|  | C | Dispersal through upper reaches of the MDB (across the GDR) | Unmack *et al.* 2001 |  | 25 |
|  | D | Two separate migration events into the MDB (via Lower and Mid) | N/A |  | 24 |
|  | E | Vicariance following the patterns of the phylogenetic tree (i.e. MID most basal) | N/A |  | 25 |
| 4 | A | Sea-level rise sequentially disconnecting river systems along coast | Williams *et al.* 2018 | T_BASE <_ 25,000 years ago  Ts = 100 – 5 ,000 years apart | 19 |
|  | B | Sea-level rise sequentially disconnecting river systems along coast, removing migration | Williams *et al.* 2018 | T_BASE <_ 25 Kya  Ts = 100 – 5,000 years apart  M_1_ – M_12_ = 10^-10^ – 10^-5^ | 29 |
|  | C | Reduction of Lake Corangamite isolating eastern populations along the coast | White, 2000 | T_BAR_ = 1,000 years ago  M_1_ – M_12_ = 10^-10^ – 10^-5^ | 18 |
|  | D | Reduction of Lake Corangamite isolating eastern populations along the coast, removing migration | White, 2000 | T_BAR_ = 1,000 years ago | 26 |
|  | E | Both sea-level rise and reduction in Lake Corangamite contributing to disconnection of river systems along coast, removing migration | White, 2000 | T_BAR_ = 1,000 years ago  T_BASE <_ 25,000 years ago  Ts = 100 – 5 ,000 years apart | 31 |
| 5 | A | Divergence between MID and the rest of the MDB populations prior to European settlement | Cole *et al.* 2016 | T_DIV_ > 200 years ago | 4 |
|  | B | Divergence between MID and the rest of the MDB populations after European settlement | Brauer *et al.* 2016 | T_DIV_ < 200 years ago | 4 |
|  | C | Divergence between MID and the rest of the MDB populations prior to European settlement with post-isolation gene flow | Cole *et al.* 2016 | T_DIV_ > 200 years ago | 6 |
|  | D | Divergence between Upper and the Lower MDB populations prior to European settlement | Cole *et al.* 2016 | T_DIV_ > 200 years ago | 4 |
|  | E | Divergence between Upper and the Lower MDB populations after European settlement | Brauer *et al.* 2016 | T_DIV_ < 200 years ago | 4 |
| 6 | A | Divergence of populations of *N.* ‘flindersi’ across the Bass Strait due to vicariance from early cycles of marine inundation | Coleman *et al.* 2010 | T_TAS_ = 1.5 – 2 Mya  T_VIC_ = 2 – 2.5 Mya | 6 |
|  | B | Divergence of populations of *N.* ‘flindersi’ across the Bass Strait due to variance from early cycles of marine inundation with gene flow post-isolation | Coleman *et al.* 2010 | M_1_ – M_6_ = 10^-10^ – 10^-5^  T_TAS_ = 1.5 – 2 Mya  T_VIC_ = 2 – 2.5 Mya | 10 |
|  | C | Divergence of populations of *N. ‘*flindersi’ across the Bass Strait due to isolated dispersal events with associated bottleneck | N/A | T_TAS_ = 1.5 – 2 Mya  T_VIC_ = 2 – 2.5 Mya  R_SRLO_ = -10^-4^ – 0  R_ART_ = -10^-4^ – 0  R_FI_ = -10^-4^ – 0 | 7 |
|  | D | Divergence of populations of *N. ‘*flindersi’ across the Bass Strait due to isolated dispersal events with associated bottleneck and gene flow post-isolation | N/A | M_1_ – M_6_ = 10^-10^ – 10^-5^  T_TAS_ = 1.5 – 2 Mya  T_VIC_ = 2 – 2.5 Mya  R_SRLO_ = -10^-4^ – 0  R_ART_ = -10^-4^ – 0  R_FI_ = -10^-4^ – 0 | 11 |

| **Model** | **Total number of parameters** | **∆Likelihood** | **AIC** | **∆AIC** | **Akaike Weight** |
| --- | --- | --- | --- | --- | --- |
| 1A | 4 | 939.081 | 38816.7869 | 1915.140203 | 0 |
| **1B** | **6** | **522.345** | **36901.6467** | **0** | **1** |
| 1C | 6 | 6368.01 | 63821.92887 | 26920.28218 | 0 |
| 1D | 8 | 719.992 | 37815.84477 | 914.1980717 | 3.0508 x 10^-199^ |
| 2A | 7 | 22822.537 | 228207.3438 | 91919.23483 | 0 |
| **2B** | **13** | **2859.923** | **136288.1089** | **0** | **1** |
| 2C | 8 | 25908.698 | 244537.9509 | 108249.842 | 0 |
| 2D | 10 | 22404.886 | 228406.3004 | 92118.19146 | 0 |
| 2E | 16 | 3777.704 | 142636.9572 | 6348.848263 | 0 |
| **3A** | **25** | **30794.197** | **914430.059** | **0** | **1** |
| 3B | 25 | 58640.469 | 1042666.881 | 128236.8216 | 0 |
| 3C | 25 | 58671.987 | 1042812.026 | 128381.9674 | 0 |
| 3D | 25 | 104816.136 | 1284161.485 | 369731.4258 | 0 |
| 3E | 25 | 36531.741 | 940852.4256 | 26422.36657 | 0 |
| 4A | 19 | 4427.185 | 234356.2201 | 3570.03817 | 0 |
| 4B | 27 | 4456.823 | 234508.7081 | 3722.526204 | 0 |
| 4C | 18 | 4298.586 | 233761.9998 | 2975.817889 | 0 |
| 4D | 26 | 4218.219 | 233407.8961 | 2621.714177 | 0 |
| **4E** | **26** | **3648.921** | **230786.1819** | **0** | **1** |
| 5A | 4 | 520.131 | 14400.71798 | 221.3401875 | 8.6415 x 10^-49^ |
| 5B | 4 | 531.54 | 14453.25837 | 273.8805742 | 3.36968 x 10^-60^ |
| **5C** | **6** | **471.199** | **14179.3778** | **0** | **1** |
| 5D | 4 | N/A | N/A | N/A | N/A |
| 5E | 4 | N/A | N/A | N/A | N/A |
| 6A | 7 | 3463.211 | 36803.27241 | 10847.18472 | 0 |
| **6B** | **13** | **1101.337** | **25938.44068** | **0** | **1** |
| 6C | 10 | 3561.24 | 37260.71264 | 11304.62494 | 0 |
| 6D | 16 | 1129.353 | 26073.45913 | 117.3714358 | 4.79862 x 10^-30^ |

**Table S3:** Coalescent model likelihoods based on biogeographic hypotheses using FastSimCoal2. Individual model specifications and their underlying hypotheses are detailed in Table S2 and Figure S5. Model likelihoods are reported as the difference between the maximum estimated likelihood and the observed likelihood of the SFS under each model. For each set of models, comparative likelihoods were calculated using Akaike Information Criterion (AIC) as [(2 x number of parameters) – (2 x log likelihood of the model)]. The difference between the AIC of each model and the lowest AIC of the model set (∆AIC) was also used to evaluate comparative likelihoods. Models were also compared using Akaike Weights to better characterise the fit of a model within a set. N/A denotes models which could not be estimated even under broad priors due to non-coalescence of demes. The best model of each set is indicated in bold.

**Table S4:** Pearson’s pairwise correlation for all 19 bioclimatic variables obtained from WorldClim v1.4. Highly correlated variables, either positively or negatively (R ≥ 0.8) are highlighted in bold.

| **Variable** | **Description** | **Bio1** | **Bio2** | **Bio3** | **Bio4** | **Bio5** | **Bio6** | **Bio7** | **Bio8** | **Bio9** | **Bio10** | **Bio11** | **Bio12** | **Bio13** | **Bio14** | **Bio15** | **Bio16** | **Bio17** | **Bio18** | **Bio19** |
| --- | --- | --- | --- | --- | --- | --- | --- | --- | --- | --- | --- | --- | --- | --- | --- | --- | --- | --- | --- | --- |
| **Bio1** | Annual Mean Temp. |  |  |  |  |  |  |  |  |  |  |  |  |  |  |  |  |  |  |  |
| **Bio2** | Mean Diurnal Range (Mean of monthly (max temp - min temp)) | 0.627 |  |  |  |  |  |  |  |  |  |  |  |  |  |  |  |  |  |  |
| **Bio3** | Isothermality (x100) | -0.062 | -0.155 |  |  |  |  |  |  |  |  |  |  |  |  |  |  |  |  |  |
| **Bio4** | Temp. Seasonality (std. dev. x100) | 0.563 | **0.856** | -0.627 |  |  |  |  |  |  |  |  |  |  |  |  |  |  |  |  |
| **Bio5** | Max Temp. Warmest Month | **0.897** | **0.882** | -0.245 | **0.834** |  |  |  |  |  |  |  |  |  |  |  |  |  |  |  |
| **Bio6** | Min Temp. Coldest Month | 0.626 | -0.123 | 0.404 | -0.263 | **0.877** |  |  |  |  |  |  |  |  |  |  |  |  |  |  |
| **Bio7** | Temp. Annual Range | 0.593 | **0.951** | -0.448 | **0.972** | **0.877** | -0.232 |  |  |  |  |  |  |  |  |  |  |  |  |  |
| **Bio8** | Mean Temp. Wettest ¼ | 0.675 | 0.470 | -0.273 | 0.547 | 0.618 | 0.210 | 0.519 |  |  |  |  |  |  |  |  |  |  |  |  |
| **Bio9** | Mean Temp. Driest ¼ | 0.310 | 0.176 | 0.223 | 0.018 | 0.287 | 0.405 | 0.087 | -0.349 |  |  |  |  |  |  |  |  |  |  |  |
| **Bio10** | Mean Temp. Warmest ¼ | **0.958** | 0.774 | -0.266 | 0.775 | **0.975** | 0.391 | 0.788 | 0.700 | 0.249 |  |  |  |  |  |  |  |  |  |  |
| **Bio11** | Mean Temp. Coldest ¼ | **0.893** | 0.289 | 0.267 | 0.131 | 0.626 | **0.899** | 0.184 | 0.498 | 0.378 | 0.728 |  |  |  |  |  |  |  |  |  |
| **Bio12** | Annual Precip. | **-0.829** | -0.675 | -0.090 | -0.501 | **-0.804** | -0.458 | -0.583 | -0.502 | -0.337 | **-0.805** | -0.722 |  |  |  |  |  |  |  |  |
| **Bio13** | Precip. Wettest Month | **-0.805** | -0.723 | -0.023 | -0.568 | **-0.817** | -0.357 | -0.646 | -0.522 | -0.304 | **-0.809** | -0.655 | 0.979 |  |  |  |  |  |  |  |
| **Bio14** | Precip. Driest Month | -0.786 | -0.504 | -0.263 | -0.282 | -0.688 | -0.641 | -0.375 | -0.346 | -0.435 | -0.698 | -0.794 | 0.911 | **0.825** |  |  |  |  |  |  |
| **Bio15** | Precip. Seasonality (Coefficient of Variation) | **-0.811** | -0.404 | 0.512 | -0.557 | -0.287 | 0.468 | -0.523 | -0.344 | 0.222 | -0.257 | 0.209 | 0.098 | 0.268 | -0.254 |  |  |  |  |  |
| **Bio16** | Precip. Wettest ¼ | **-0.811** | -0.725 | -0.010 | -0.578 | **-0.822** | -0.354 | -0.652 | -0.543 | -0.283 | **-0.817** | -0.656 | 0.982 | **0.998** | **0.825** | 0.268 |  |  |  |  |
| **Bio17** | Precip. Driest ¼ | -0.782 | -0.524 | -0.256 | -0.695 | -0.695 | -0.618 | -0.393 | -0.354 | -0.424 | -0.701 | -0.781 | 0.936 | **0.857** | **0.991** | -0.214 | **0.858** |  |  |  |
| **Bio18** | Precip. Warmest ¼ | -0.722 | -0.457 | -0.292 | -0.223 | -0.634 | -0.639 | -0.322 | -0.206 | -0.540 | -0.631 | -0.753 | 0.880 | **0.802** | **0.966** | -0.243 | 0.797 | **0.966** |  |  |
| **Bio19** | Precip. Coldest ¼ | -0.799 | -0.734 | 0.036 | -0.609 | **-0.818** | -0.301 | -0.675 | -0.608 | -0.202 | **-0.818** | -0.624 | 0.961 | **0.986** | 0.775 | 0.323 | **0.990** | **0.812** | 0.725 |  |

**Table S5:** Pearson’s pairwise correlation for 9 uncorrelated bioclimatic variables obtained from Worldclim v1.4, used in estimating species and lineage distribution models in MaxEnt.

| **Variable** | **Description of variable** | **Bio1** | **Bio2** | **Bio3** | **Bio6** | **Bio8** | **Bio9** | **Bio14** | **Bio15** | **Bio19** |
| --- | --- | --- | --- | --- | --- | --- | --- | --- | --- | --- |
| **Bio1** | Annual Mean Temperature |  |  |  |  |  |  |  |  |  |
| **Bio2** | Mean Diurnal Range (Mean of monthly (max temp - min temp)) | 0.63 |  |  |  |  |  |  |  |  |
| **Bio3** | Isothermality (BIO2/BIO7) (* 100) | -0.06 | -0.16 |  |  |  |  |  |  |  |
| **Bio6** | Min Temperature of Coldest Month | 0.63 | -0.12 | 0.40 |  |  |  |  |  |  |
| **Bio8** | Mean Temperature of Wettest Quarter | 0.67 | 0.47 | -0.27 | 0.21 |  |  |  |  |  |
| **Bio9** | Mean Temperature of Driest Quarter | 0.31 | 0.18 | 0.22 | 0.41 | -0.35 |  |  |  |  |
| **Bio14** | Precipitation of Driest Month | -0.79 | -0.50 | -0.26 | -0.64 | -0.35 | -0.44 |  |  |  |
| **Bio15** | Precipitation Seasonality (Coefficient of Variation) | -0.08 | -0.40 | 0.51 | 0.47 | -0.34 | 0.22 | -0.25 |  |  |
| **Bio19** | Precipitation of Coldest Quarter | -0.80 | -0.73 | 0.04 | -0.30 | -0.61 | -0.20 | 0.78 | 0.32 |  |

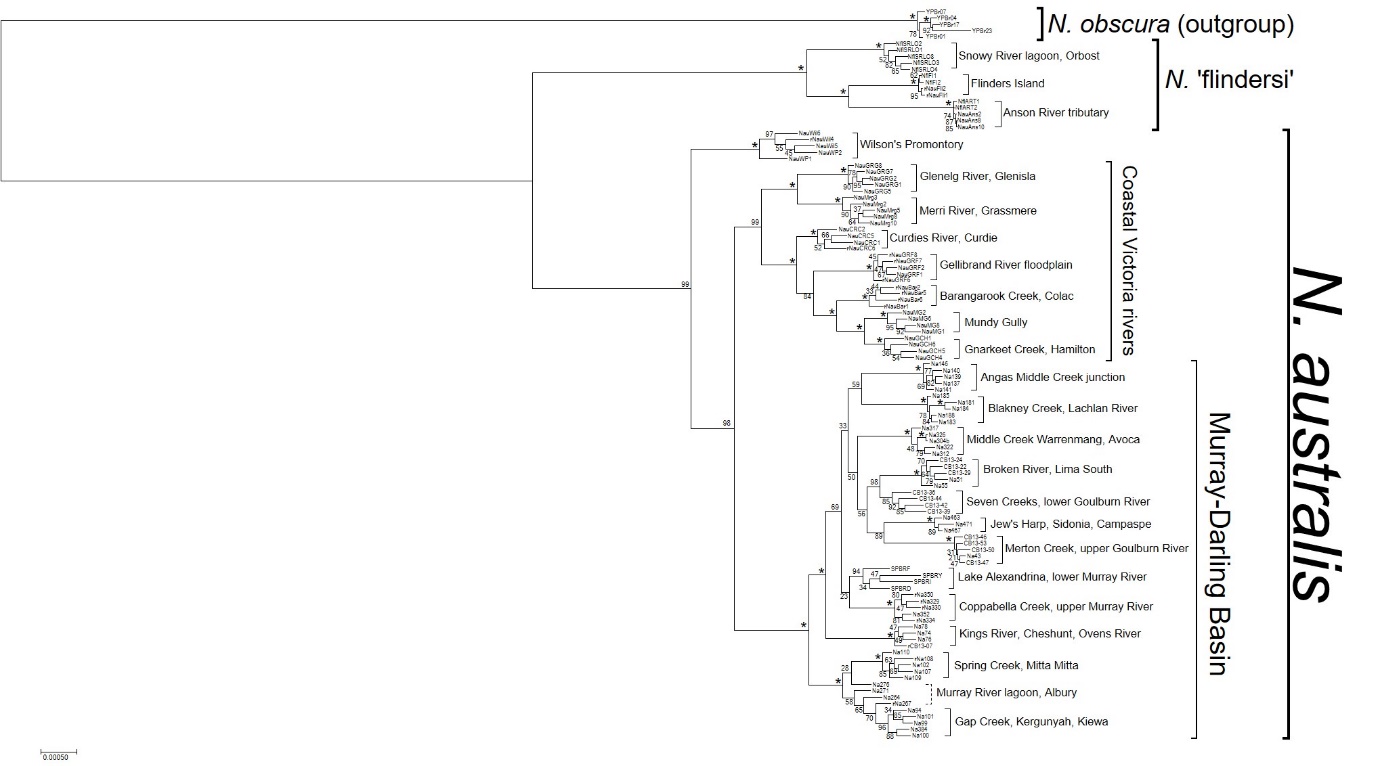

**Figure S1:** Maximum likelihood phylogeny of all 114 *N. australis* and *N.* ‘flindersi’ samples based on 7,780 concatenated RADseq loci. Node labels indicate bootstrap support from 1,000 RELL bootstraps, with asterisks denoting nodes with 100% support. Species, clades and populations are denoted by square brackets.

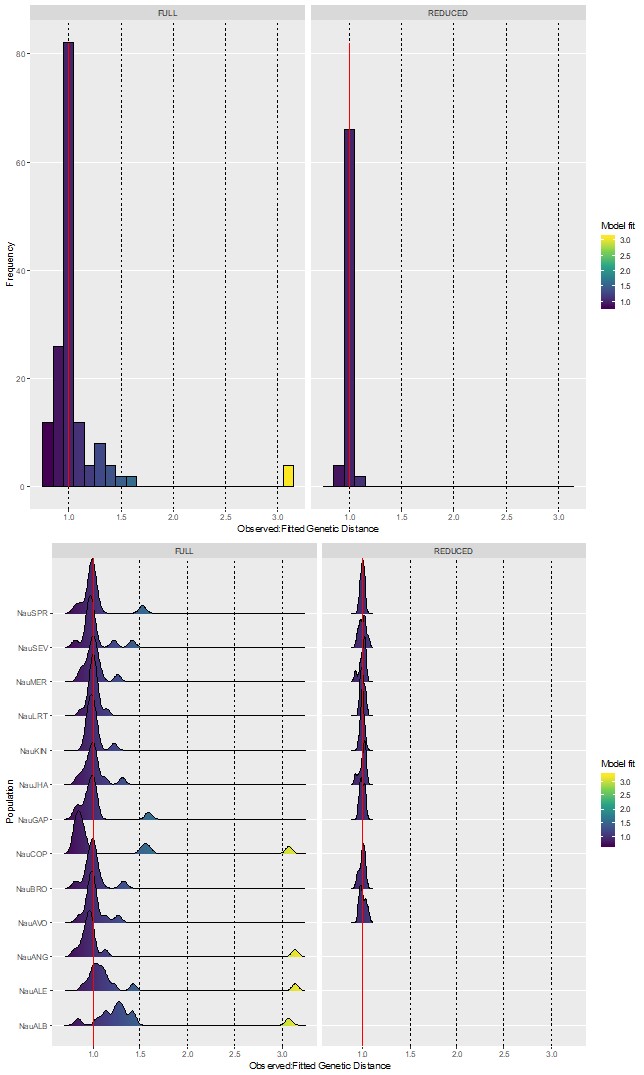

**Figure S2:** Evaluation of *StreamTree* model fit across the entire set of population comparisons (top) and for comparisons involving each specific population (bottom). Density plots represent the probability distribution of mismatch (ratio) between observed (true) genetic distance and distance modelled by *StreamTree*, with the red line indicating perfect fit of the model (StreamTree value = observed genetic distance). The full dataset (left) is compared with the model with divergent outliers removed (right).

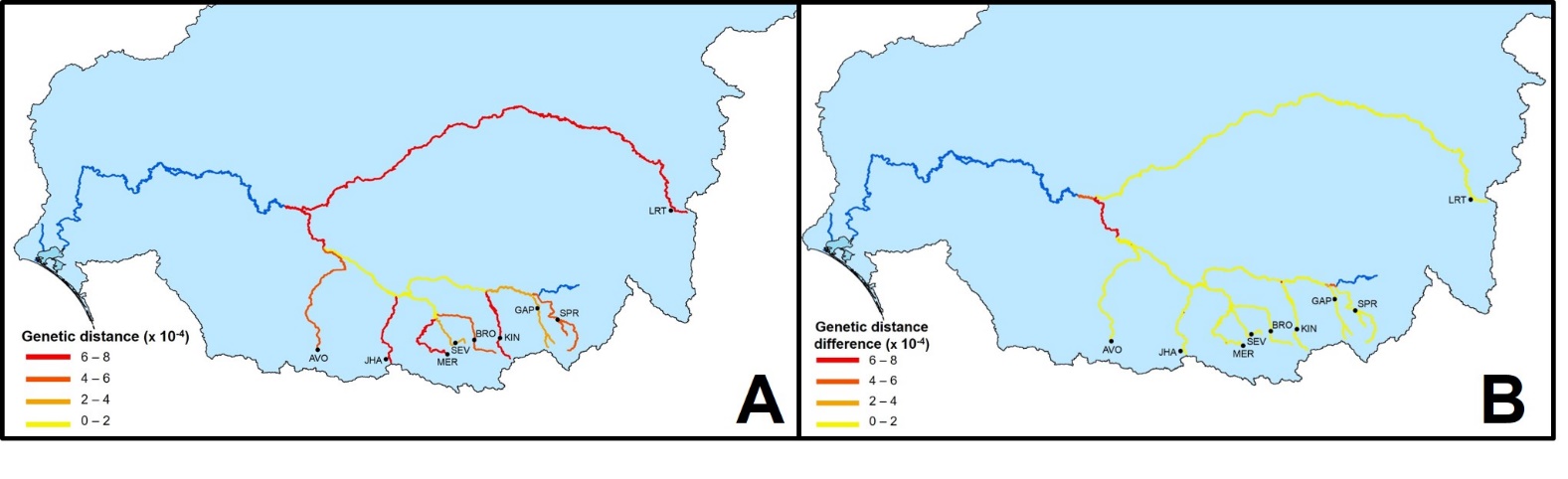

**Figure S3:** Visual representation of the reduced *StreamTree* model (excluding 4 outlier populations) of genetic divergence across the Murray-Darling Basin (MDB). **A:** Dendritic riverine network of the MDB, with streams colour-coded according to the reduced *StreamTree* model that determines the contribution (as a penalty) of each segment in driving genetic divergence (measured as mean uncorrected genetic distance per population (*p*) x 10^2^) across the basin. Segments coloured in yellow confer little penalty (i.e. genetic divergence between populations at either end of the segment is low) whereas red segments confer higher genetic differentiation. **B:** Visual comparison of *StreamTree* models using the full (*n* = 13 populations) and reduced (*n* = 9) populations for stream segments that were considered under both models. Yellow segments demonstrate streams with similar associated penalties across both models whereas red segments showed more variable penalties between the two models.

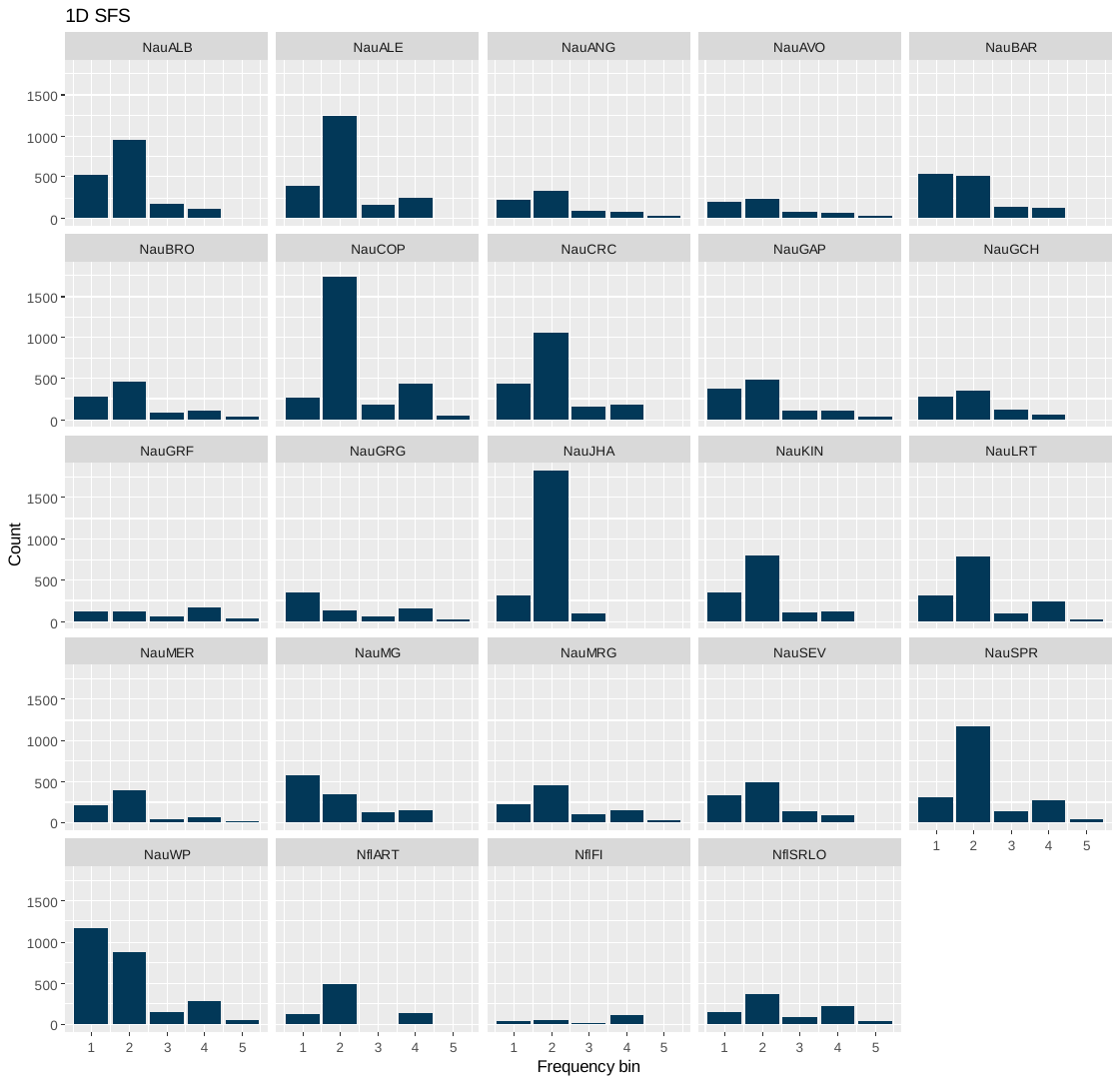

**Figure S4:** One-dimensional site-frequency spectra for stairway plot analysis. SNPs were called independently for each population and hence are variable in number across the populations.

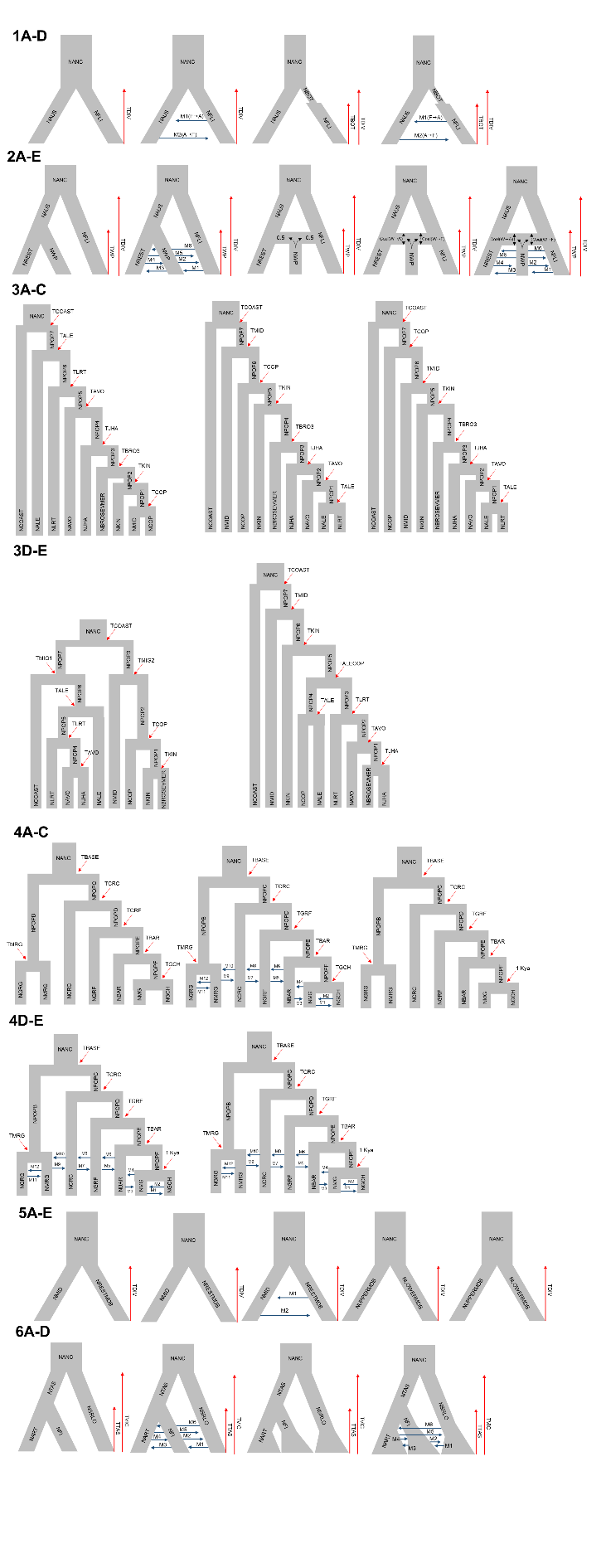

**Figure S5:** Diagrammatic representations of coalescent models tested within FastSimCoal2.6, as described in Table S2. Red arrows denote divergence time parameters whilst blue arrows denote migration rate parameters (in terms of gene flow forward in time in the direction of the arrow).

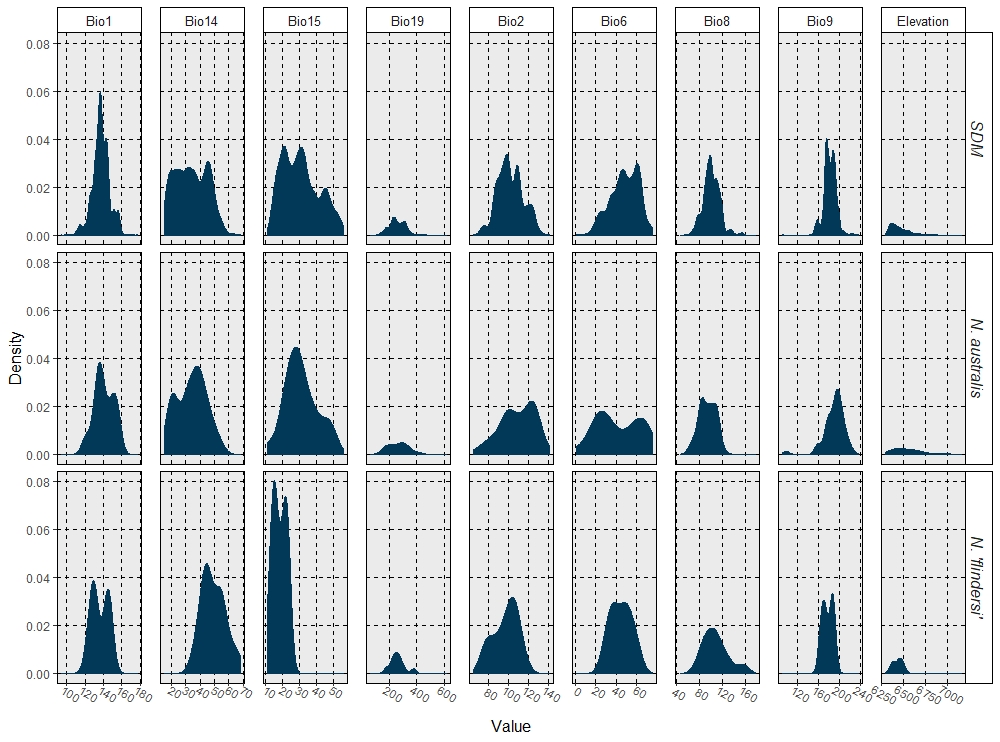

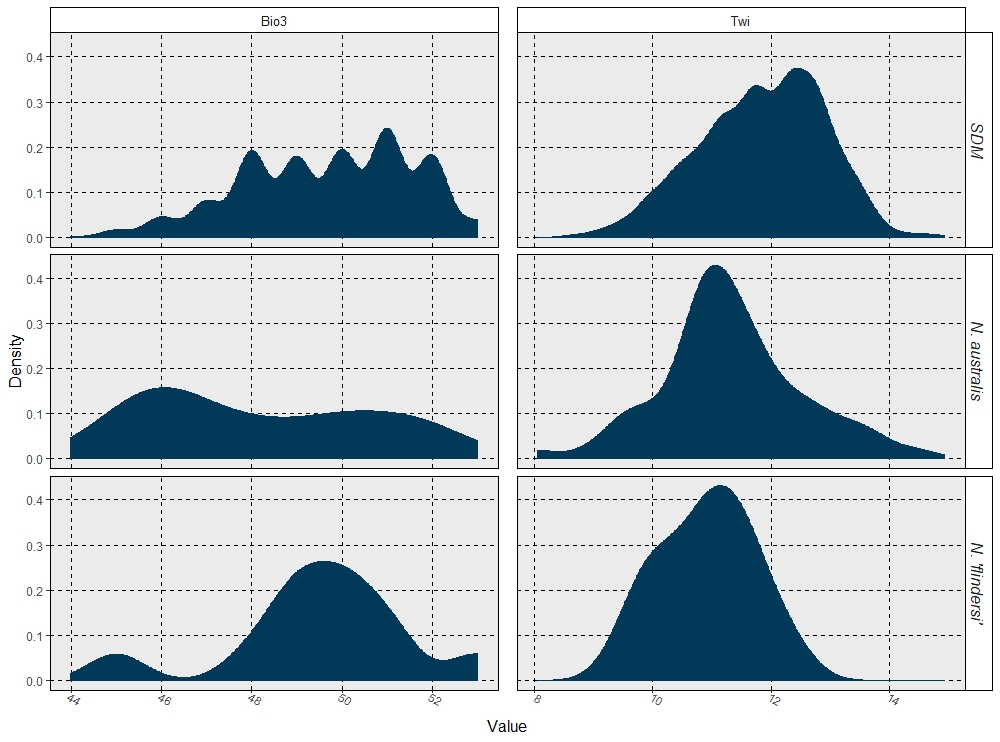

**Figure S6:** Relative density plots of environmental variables within the total species distribution model (SDM) and each lineage-distribution model (LDM) individually. Variable Bio3 and Twi are plotted separately due to variance in maximum density. Environmental data across all variables were sampled from points using ArcMap based on 2,528 SDM occurrences, 61 *N. australis* LDM occurrences and 11 *N.* ‘flindersi’ LDM occurrences.
